# Synthesizing Images of Tau Pathology from Cross-modal Neuroimaging using Deep Learning

**DOI:** 10.1101/2022.09.07.507042

**Authors:** Jeyeon Lee, Brian J. Burkett, Hoon-Ki Min, Matthew L. Senjem, Ellen Dicks, Nick Corriveau-Lecavalier, Carly T. Mester, Heather J. Wiste, Emily S. Lundt, Melissa E. Murray, Aivi T. Nguyen, Ross R. Reichard, Hugo Botha, Jonathan Graff-Radford, Leland R. Barnard, Jeffrey L. Gunter, Christopher G. Schwarz, Kejal Kantarci, David S. Knopman, Bradley F. Boeve, Val J. Lowe, Ronald C. Petersen, Clifford R. Jack, David T. Jones

## Abstract

Given the prevalence of dementia and the development of pathology-specific disease modifying therapies, high-value biomarker strategies to inform medical decision making are critical. In-vivo tau positron emission tomography (PET) is an ideal target as a biomarker for Alzheimer’s disease diagnosis and treatment outcome measure. However, tau PET is not currently widely accessible to patients compared to other neuroimaging methods. In this study, we present a convolutional neural network (CNN) model that impute tau PET images from more widely-available cross-modality imaging inputs. Participants (n=1,192) with brain MRI, fluorodeoxyglucose (FDG) PET, amyloid PET, and tau PET were included. We found that a CNN model can impute tau PET images with high accuracy, the highest being for the FDG-based model followed by amyloid PET and MRI. In testing implications of AI-imputed tau PET, only the FDG-based model showed a significant improvement of performance in classifying tau positivity and diagnostic groups compared to the original input data, suggesting that application of the model could enhance the utility of the metabolic images. The interpretability experiment revealed that the FDG- and MRI-based models utilized the non-local input from physically remote ROIs to estimate the tau PET, but this was not the case for the PiB-based model. This implies that the model can learn the distinct biological relationship between FDG PET, MRI, and tau PET from the relationship between amyloid PET and tau PET. Our study suggests that extending neuroimaging’s use with artificial intelligence to predict protein specific pathologies has great potential to inform emerging care models.

## INTRODUCTION

Misfolded tau neurofibrillary tangles (NFT) are the characteristic pathologic feature of tauopathies, a group of progressive neurodegenerative disease entities. Together with amyloid-β plaques, tau NFTs are the classic pathologic feature of the most common etiology of dementia, Alzheimer’s disease (AD).^1,2^ Tau NFT burden is important because it correlates with the degree of cognitive impairment in AD and neuropathologic studies support a stronger correlation of tau pathology than amyloid-β plaques to cognitive status.^3–5^

Tau positron emission tomography (PET), which is a minimal invasive method to quantify the extent and distribution of pathologic NFT in the brain,^6–8^ is therefore a promising tool to assess response to therapy or changes over time.^9^ Cross-sectional studies show that tau PET uptake levels can be used effectively to support a clinical diagnosis of AD dementia and to estimate disease severity.^10–14^ Tau PET uptake patterns *in vitro* have been associated with Braak tangle stage.^8,15^ Other studies show that the tau PET signal is associated with aging^16,17^ and with reduced glucose metabolism.^11,18,19^ Furthermore, tau PET uptake patterns have been correlated to specific clinical phenotypes of AD, whereas amyloid PET has not, with distinct distributions of tau pathology associated with posterior cortical atrophy, logopenic variant primary progressive aphasia, and other presentations of AD.^11,20–24^ PET studies of tau accumulation show a close spatial relationship with gray matter volume reduction in clinical variants of AD.^11,20,25,26^

Accordingly, clinical interest in tau PET has grown as a tool to measure and visualize in vivo tau pathology, for both AD and other tauopathies. Multiple tau PET agents have been developed for this purpose.^7^ Recently, [^18^F]flortaucipir ([^18^F]AV-1451) received FDA approval for clinical use in the evaluation of AD.^27^ This ligand has been shown to have specificity for AD-like tau pathology in vivo^28^ and has been used to stratify participants included in a recent clinical trial targeting amyloid pathology.^29^ However, at present, tau PET is not widely accessible to patients compared to other neuroimaging methods.^30^ Moreover, the addition of tau PET to the diagnostic evaluation of dementia, which currently includes [^18^F]fluorodeoxyglucose (FDG) PET and amyloid PET, creates an additional burden on patients of undergoing the test and the exposure to multiple radiopharmaceuticals. In addition, FDG-PET has wide application across all forms of degenerative dementia because it contains useful features across the entire spectrum of etiologies beyond amyloid and tau associated conditions.^31^ Nevertheless, measuring tau pathology is integral to the diagnosis and prognosis of the AD continuum, and increasing the accessibility of tau PET has potential to enable a greater role in research and clinical applications in the future.^32^

In this study, we developed a convolutional neural network (CNN) model which enables a cross-modal tau PET synthesis using other neuroimaging data, including FDG PET, structural T1-weighted magnetic resonance imaging (MRI) or amyloid PET, as input. A relationship between tau PET and other modalities has been proposed and tau burden has been correlated to regions of FDG hypometabolism,^11,18,22,33^ cortical atrophy,^13,20,34,35^ and amyloid accumulation.^34,36–38^ Therefore, in the absence of tau PET, it is plausible that valid information regarding tau pathology may be inferred from other neuroimaging data, although the relationship is complex. We hypothesized that the CNN model trained on a large neuroimaging sample might enable an accurate estimation of tau PET images by learning the underlying biological relationship between biomarkers. With recent advances in deep learning techniques, several works have explored cross-modality synthesis that transforms images from one domain to another, including low-dose FDG PET to standard-dose FDG PET,^39^ computed tomography (CT) to MRI,^40^ MRI to FDG PET,^41,42^ and CT to FDG PET.^43^ In the current work, we present a 3D dense-U-net model for the imputation of tau PET from either FDG PET, amyloid PET, or structural MRI, evaluating the performance of the AI-imputed tau PET data using ground-truth tau PET.

## RESULTS

### Participants and Data

Participants who underwent MRI, FDG PET, amyloid PET with Pittsburgh compound B (PiB)^44^ and tau PET with ^18^F-flortaucipir (AV-1451)^45^ were included (n=1,192, number of scans=1,505)(Table 1; see Fig.S1 for data inclusion/exclusion criteria). The participants were categorized into major clinical subgroups based on clinical diagnosis including cognitively unimpaired (CU; n=739), mild cognitive impairment (MCI; n=169), typical AD (n=110), behavioral variant of frontotemporal dementia (bvFTD; n=25), dementia with Lewy bodies (DLB; n=38) and other clinical syndromes (e.g., vascular cognitive impairment, idiopathic REM sleep behavior disorder (RBD), posterior cortical atrophy (PCA), semantic dementia (SD), logogenic variant of primary progressive aphasia (lvPPA), non-fluent variant of primary progressive aphasia (nfvPPA) and progressive supranuclear palsy (PSP); n=111) (Table 1). The clinical categories in these databases were not used in training the algorithm, given that ground-truth was the tau PET scan from these participants. However, we evaluated the implications of the trained models using common clinical categories (CU, AD-spectrum, FTD-spectrum, and DLB-spectrum).

**Table 1.** Demographics for Mayo participants. {}Brackets in the characteristic column indicate the number of participants missing this particular variable.

| Characteristic | Clinical Diagnosis |  |  |  |  |  |
| --- | --- | --- | --- | --- | --- | --- |
|  | Normal | MCI | AD | FTD | DLB | Others |
| N (%) | 739 (62) | 169 (14.18) | 110 (9.23) | 25 (2.10) | 38 (3.19) | 111 (9.31) |
| Age, median (min max), years | 69 (30 94) | 74 (26 98) | 72 (53 92) | 62 (43 75) | 70 (45 89) | 68 (33 85) |
| Male sex, n (%) | 385 (52.10) | 118 (69.82) | 53 (48.18) | 11 (44) | 33 (86.84) | 64 (57.66) |
| Education, median (IQR), years<br>{3} | 16 (13 17) | 16 (12 18) | 16 (13 17) | 16 (13.75<br>18) | 15.5 (14<br>18) | 16 (14 18) |
| Clinical Dementia Rating<br>Scale-Sum of Boxes, median<br>(IQR) {9} | 0 (0 0) | 1 (0.5 1.5) | 4.5 (2.5 7) | 4.25 (2.75<br>6.25) | 4.5 (3 6) | 1.5 (0.5 4) |
| Meta-ROI FDG PET SUVR,<br>median (IQR) | 1.55<br>(1.46 1.64) | 1.39<br>(1.28 1.49) | 1.19<br>(1.05 1.30) | 1.42<br>(1.32 1.55) | 1.17<br>(1.08 1.27) | 1.42<br>(1.22 1.56) |
| Meta-ROI PIB PET SUVR,<br>median (IQR) {19} | 1.39<br>(1.31 1.52) | 1.55<br>(1.37 2.32) | 2.49<br>(2.23 2.72) | 1.34<br>(1.20 1.46) | 1.61<br>(1.36 2.22) | 1.43<br>(1.31 2.14) |
| Meta-ROI Tau PET SUVR,<br>median (IQR) | 1.18<br>(1.13 1.23) | 1.24<br>(1.19 1.40) | 1.91<br>(1.60 2.31) | 1.25<br>(1.18 1.35) | 1.21<br>(1.17 1.28) | 1.26<br>(1.16 1.55) |

The 3D Dense-U-Net architecture was utilized for the CNN model (Fig.1A),^46,47^ which has dense interconnections between convolutional layers (dense block). The model uses input volumes of size 128×128×128 to impute tau PET images with the same dimensions. For the image preprocessing, scans were firstly normalized to Mayo Clinic Adult Lifespan Template (MCALT) space^48^ using Unified Segmentation in SPM12.^49^ The tau- and amyloid-PET standardized uptake value ratios (SUVR) were calculated by dividing the median uptake in the cerebellar crus grey matter. FDG PET SUVR was calculated by dividing the median uptake in the pons. For T1-weighted MRIs, voxels’ intensities were normalized by dividing a mean intensity derived from individualized white matter mask after spatial normalization. Cross-validation experiments were conducted using 5-fold validations (60% training set, 20% validation set, and 20% test set) (see the method section for details). Demographics for participants in the training, validation, and testing data sets for each fold are summarized and compared in table S1 and S2 including pertinent clinical variables, measures of cognitive performance, and tau PET meta-ROI SUVR as a ground-truth measure of model performance (Fig.S2). The composition for each group was relatively similar.

**Fig. 1.**
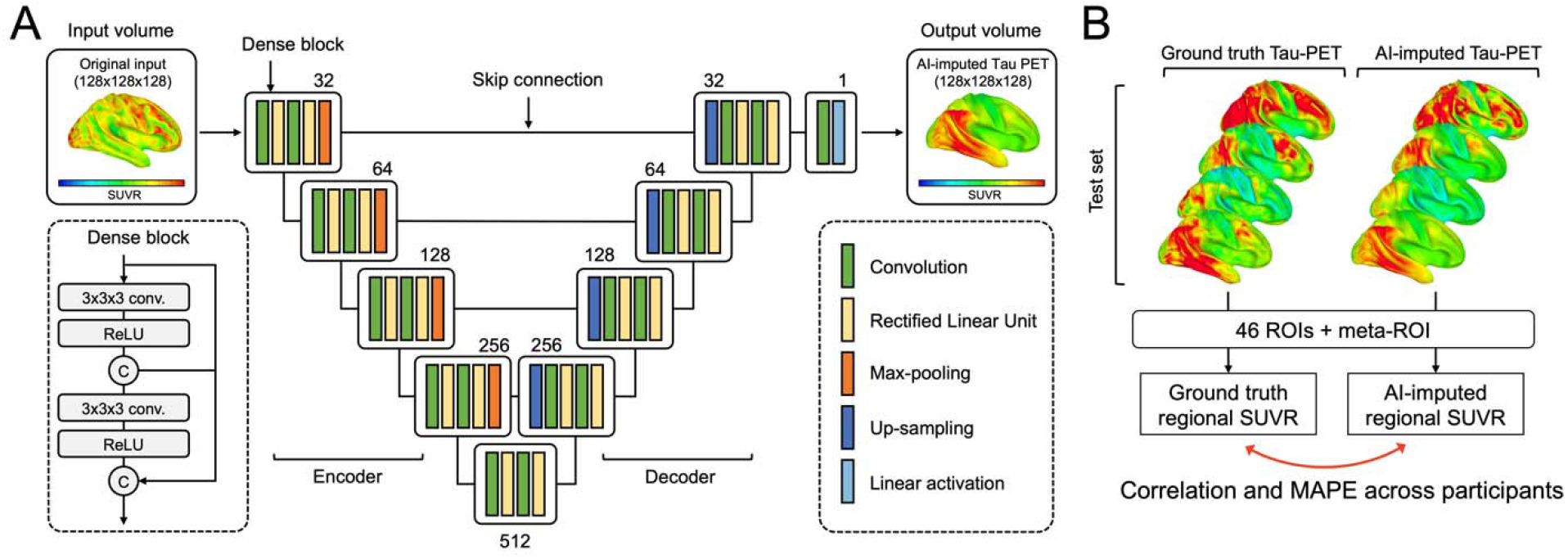
Dense-U-Net architecture and layout of analysis. **(A)** The architecture receives input of size 128×128×128 and produces the AI-imputed tau-PET of the same dimension with input data. Dense-U-net architecture is composed of encoder (left), decoder (right) and bridge. Left dotted box illustrates a layout of dense connection in dense block, when output from each rectified linear unit (ReLU) layer is concatenated (circular C) to the input of the block before fed to the next layer. The numbers denoted above the dense blocks indicate a number of filters. **(B)** The similarity between ground-truth tau PET and AI-imputed tau PET was assessed across total participants in test set using a regional SUVR calculated from 46 ROIs and meta-ROI. Pearson’s correlation and mean absolute percentage error (MAPE) were used as an evaluation metric.

### Detailed images of tau pathology in the brain can be successfully imputed from images of glucose utilization

First, we tried to impute tau PET using glucose metabolism images obtained by FDG PET. We found the dense-U-net model was able to successfully impute tau PET images from standard FDG PET. Fig.2A shows eight representative example cases from the test set, comparing the original FDG PET, ground-truth tau PET, and AI-imputed tau PET. As illustrated in Fig.2A, the AI-imputed tau PET image showed good agreement with ground-truth images in visual assessment. A high degree of similarity was observed for cases with high tau burden (Case 6, 7, and 8) and cases with subtle tau tracer activity (Case 1 and 2), demonstrating the range of tau activity the model is capable of characterizing.

**Fig. 2.**
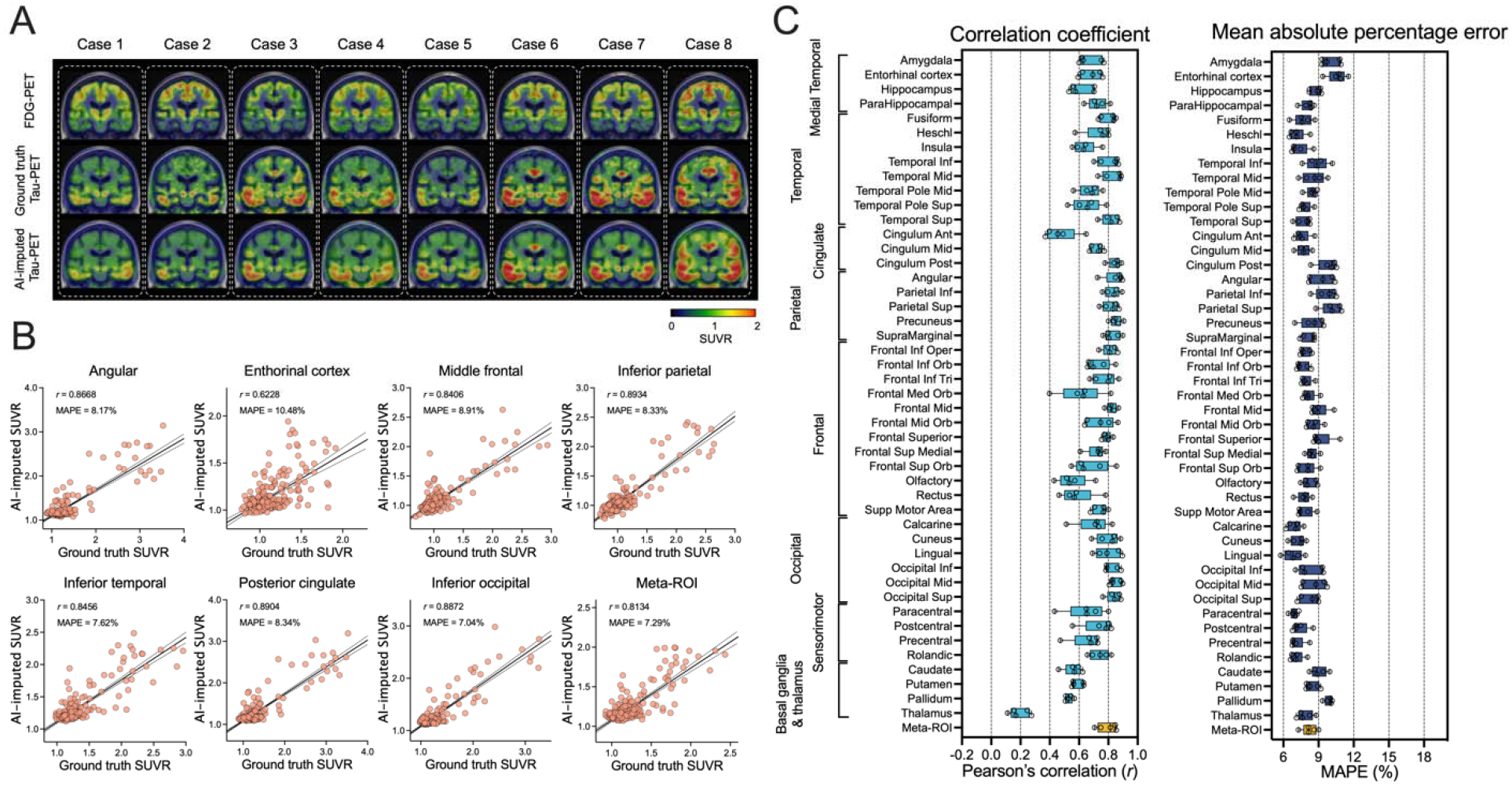
FDG-PET based tau-PET synthesis results. **(A)** Eight representative cases with original FDG-PET, ground-truth tau-PET and AI-imputed tau PET. **(B)** Scatter plots of ground-truth tau-PET and AI-imputed tau PET from seven representative ROIs and meta-ROI. r indicates Pearson’s correlation coefficient and MAPE indicates mean absolute percentage error. Linear regression (black line) and 95% confidence bands (dotted lines) are shown. **(C)** The mean of correlation coefficient and MAPE of five folds from 46 ROIs and the meta-ROI is summarized in a box plot. The yellow-colored box depicts the meta-ROI result. Open circles indicate different folds.

To quantify the model’s performance, regional SUVRs and meta-region of interest (ROI) SUVRs were extracted from both the ground-truth and AI-imputed tau PET scans (Fig.1B). The regional SUVRs were calculated by measuring median uptake in each ROI. The meta-ROI represents a set of temporal lobe ROIs and has previously been shown to have a broad dynamic range across the normal to pathological aging to AD dementia.^50^ The meta-ROI SUVR was formed from the average of the median uptake in the amygdala, entorhinal cortex, fusiform, parahippocampal and inferior temporal and middle temporal gyri.^50^ Then, the model’s performance was evaluated by Pearson’s correlation and mean absolute percentage error (MAPE) between regional SUVRs extracted from both the ground-truth and AI-imputed tau PET scans across participants. The AI-imputed tau PET SUVR, when plotted against ground-truth tau PET using a real tau tracer, demonstrated that each regional SUVR showed a high correlation (*r*>0.8) and low MAPE (~8%) as well as the meta-ROI (Fig.2B). The mean correlation coefficient and MAPE of five-fold summarized for each anatomic ROI and the meta-ROI reflect the performance of the model (Fig.2C). The mean correlation coefficient for the meta-ROI was 0.79±0.06 and the MAPE was 8.24±0.64%. The regional SUVR of the basal ganglia, a known region of off-target AV-1451 binding, showed relatively lower performance (*r*=0.53~0.58 and MAPE=8.46~9.87). The thalamic ROI showed the lowest correlation (*r*=0.19±0.07), which was part of a larger pattern where the AI-imputed images had lower predicted signal in regions of off-target binding than the actual tau PET images.

To examine whether the trained model presents a dataset-specific bias, we evaluated the performance of the Mayo-trained models on the multi-site cross-modal data from the Alzheimer’s Disease Neuroimaging initiative (ADNI) for participants that had MRI, FDG PET and tau PET (n=288; table S3). Using the ADNI scans, we observed that the regional tau PET SUVR was predicted with high accuracy from FDG PET using the dense-U-net model trained on the Mayo dataset (Fig.S3), although the overall performance slightly decreased compared to the original result from the Mayo test set (F(1,376)=386.6, p<0.001 for correlation coefficient and F(1,376)=1330, p<0.001 for MAPE, using a two-way ANOVA). For the meta-ROI, the correlation coefficient was 0.68±0.02 and the MAPE was 10.60±0.11% when derived from the ADNI dataset.

### Dense-U-net can also Impute Tau PET using Structural MRI and Amyloid PET

Next, we used the same dense-U-net architecture to impute tau PET using structural MRI. The model was initialized and separately trained from scratch. As a result, we found that the MRI-based model was also able to impute the tau PET images, although the accuracy was comparably lower than the FDG-based model (F(1,376)=424.1, p<0.001 for correlation coefficient and F(1,376)=159.5, p<0.001 for MAPE, using a twoway ANOVA; Fig.3). In some ROIs such as the insula, anterior cingulate, medial orbitofrontal, olfactory, and gyrus rectus, the correlation coefficient was considerably low (*r*<0.3), however other ROI’s regional SUVR was relatively well predicted (Fig.3B and C). In the MRI-based model, the meta-ROI’s mean correlation coefficient was 0.62±0.05 and mean MAPE was 10.16±0.82% across the 5-fold test sets.

**Fig. 3.**
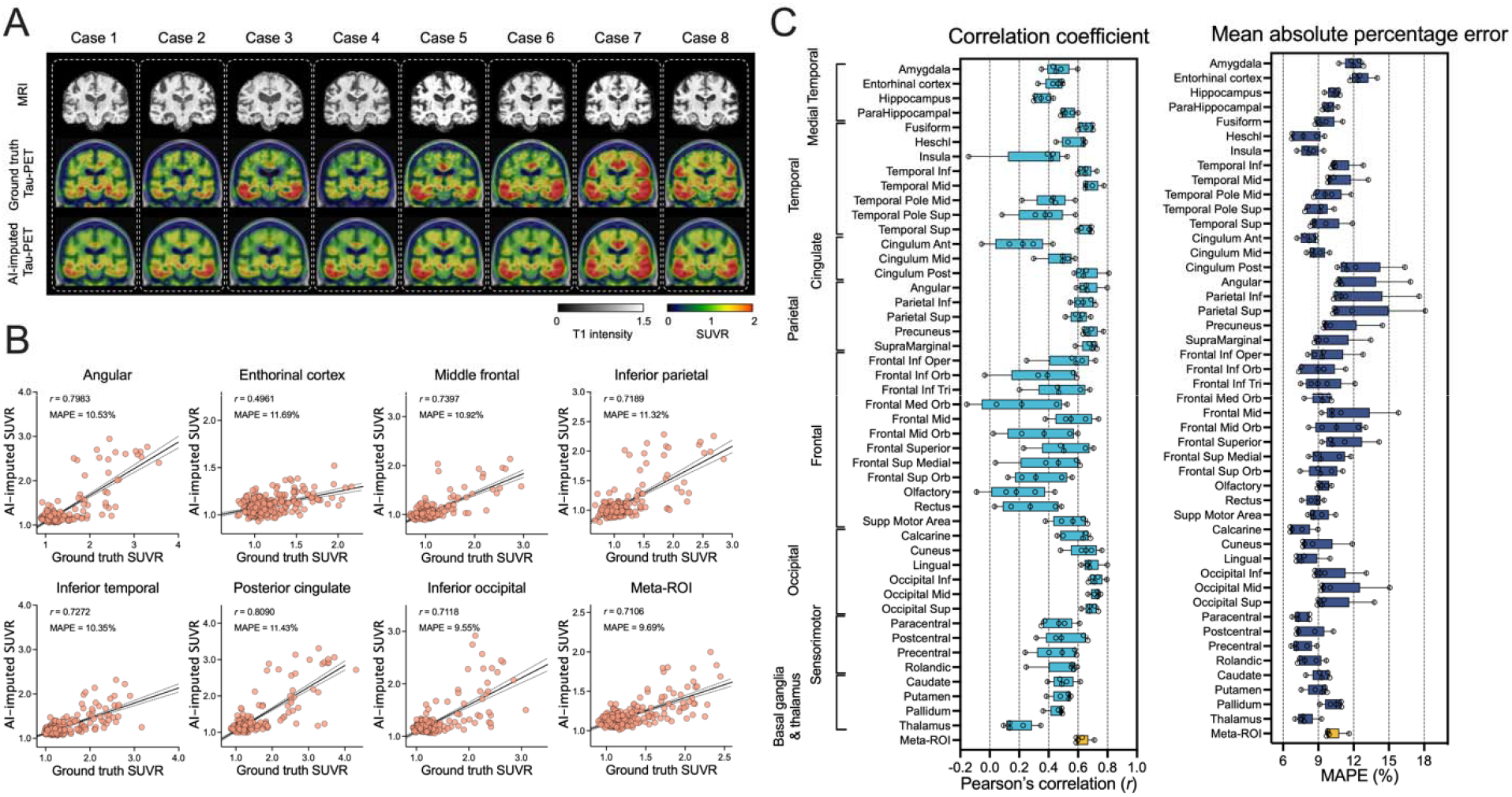
Structural MRI based tau PET synthesis results. **(A)** Eight representative cases with original MRI, ground-truth tau PET and AI-imputed tau-PET. **(B)** Scatter plots between ground-truth tau-PET and AI-imputed tau PET from seven representative ROIs and meta-ROI. r indicates the Pearson’s correlation coefficient and MAPE indicates mean absolute percentage error. Linear regression (black line) and 95% confidence bands (dotted lines) are shown. **(C)** The mean correlation coefficient and MAPE of five folds from 46 ROIs and meta-ROI is summarized in the box plots. The yellow-colored box depicts the meta-ROI result. Open circles indicate different folds.

The MRI-based model was also cross-evaluated using the ADNI dataset. As a result, the performance of the MRI-based model on an external dataset was found to have relatively low accuracy compared to the training (Mayo) dataset (F(1,376)=134.1, p<0.001 for correlation coefficient and F(1,376)=208.0, p<0.001 for MAPE, using a two-way ANOVA; Fig.S4). In most ROIs, the correlation coefficient was very low (*r*<0.2) and the MAPE was high (>10%), meaning that the MRI-based model trained on the training dataset did not show a robust performance for images acquired in a multi-site external dataset.

Next, we trained the Dense-U-net model to impute the tau PET images using amyloid PET inputs (Fig.4) from the Pittsburgh compound B (PiB) radiotracer.^44^ The PiB-based model was also able to generate AI-imputed tau PET scans (Fig.4A and B), and the mean correlation between ground-truth regional SUVR and AI-imputed regional SUVR was found to be 0.41-0.76 and the MAPE range was around 7-11% (Fig.4C). The general performance was significantly lower than the FDG-based model (F(1,376)=96.76, p<0.001 for correlation coefficient and F(1,376)=30.77, p<0.001 for MAPE, using a two-way ANOVA); however, the performance was significantly higher than the MRI-based model (F(1,376)=137.7, p<0.001 for correlation coefficient and F(1,376)=80.63, p<0.001 for MAPE, using a two-way ANOVA). The Mayo PiB-based model was not applied to the ADNI dataset because different amyloid tracers (Florbetapir^51^ and Florbetaben^52^) are used in that study.

**Fig. 4.**
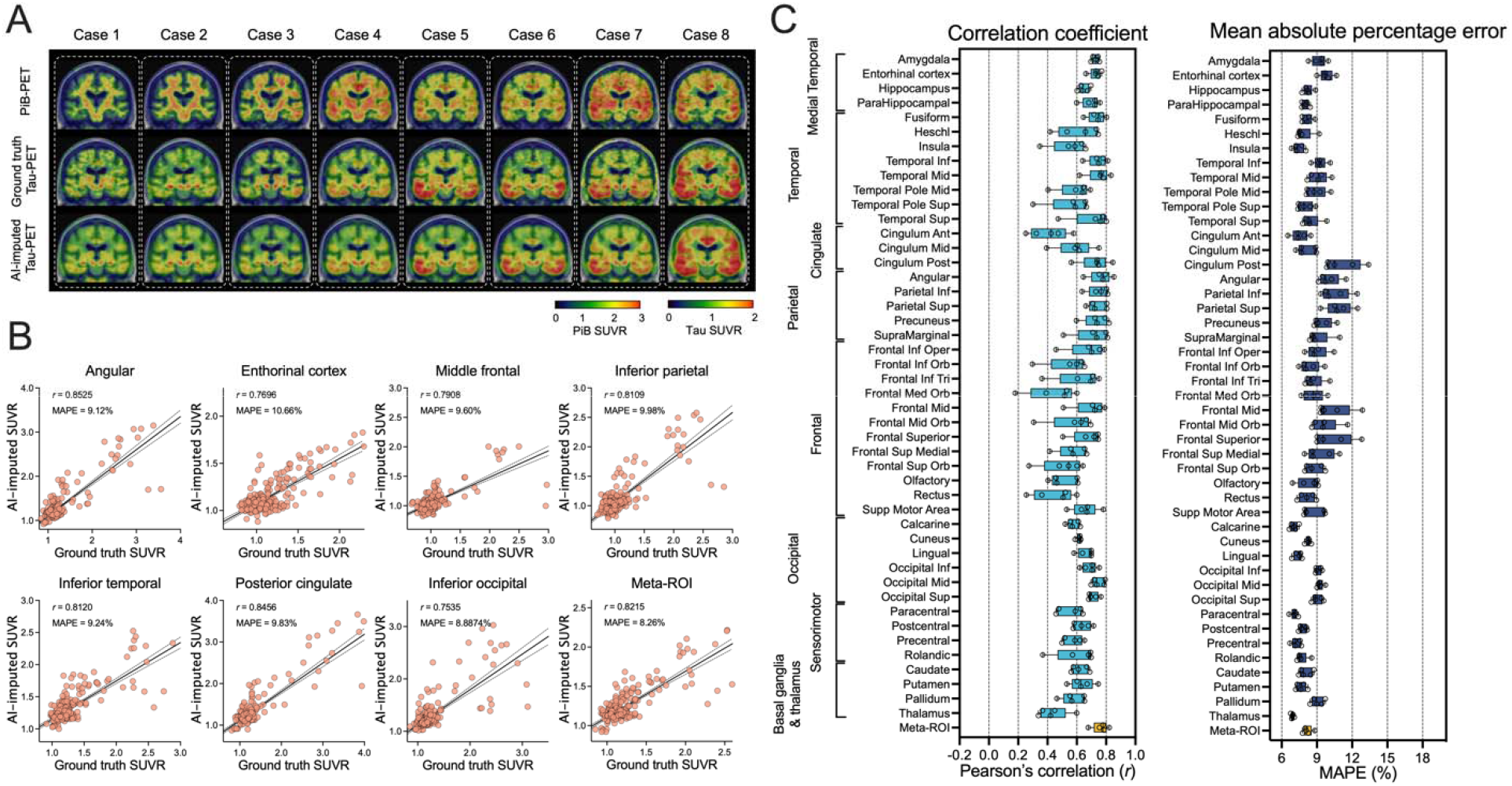
Amyloid PET based tau PET synthesis results. **(A)** Eight representative cases with actual PiB-PET, ground-truth tau PET and AI-imputed tau PET. **(B)** Scatter plots between ground-truth tau PET and AI-imputed tau PET from seven representative ROIs and meta-ROI. r indicates the Pearson’s correlation coefficient and MAPE indicates mean absolute percentage error. Linear regression (black line) and 95% confidence bands (dotted lines) are shown. **(C)** The mean of correlation coefficient and MAPE of five folds from 46 ROIs and meta-ROI is summarized in a box plot. The yellow-colored box depicts the meta-ROI result. Open circles indicate different folds.

For additional evaluation of the model’s performance, voxel-wise error maps between the AI-imputed tau and ground truth tau PET were calculated using a root mean squared error (RMSE) (Fig.S5A-C). Overall, FDG-based model showed the lowest RMSE across the cortical regions, followed by the PiB- and MRI-based model. For additional voxel-based quantitative analysis, multi-scale structural similarity index (MS-SSIM)^53^ was also computed (Fig.S5D). All modalities showed moderately high MS-SSIM values (>0.9), and the performance of both the FDG- and PiB-based models was significantly higher than the MRI-based model (p<0.001, Holm-Sidak test). Additional example images comparing ground-truth and AI-imputed tau PETs are shown in Fig.S6 and 7.

We further evaluated the accuracy of AI-imputed tau PET using postmortem neuropathology data. Thirteen participants who had tau PET within 3 years of death and complete neuropathologic assessments were eligible for evaluation. Using antibodies to phospho-tau (AT8) immunostained sections,^54,55^ Braak tangle stage^56,57^ was assessed. Then, correlations between SUVR in the meta-ROI and Braak tangle stage were calculated. For the analysis, time between tau PET and death was not specified as a covariate. As a result, the meta-ROI SUVR from the AI-imputed tau PET showed a significant correlation with the Braak stage except for the MRI-based model (p<0.05, Spearman’s correlation, Fig.S8). The correlation coefficient was not significantly different with the association between actual tau PET and Braak stage (p>0.05, z-test after Fisher’s r to z transformation, Fig.S8).

### Evaluation of Synthesized Tau PET Images Accuracy in Prediction of Tau Positivity

To assess the clinical implications of AI-imputed tau PET images, we performed ROC analyses for predicting tau positivity. In dementia research and clinical practice, although the biomarkers exist on a continuum, dichotomizing normal/abnormal tau using specific cut points is useful and widely used.^50^ We tried to predict the tau positivity obtained from the ground-truth tau PET data using four different meta-ROI cutoff thresholds (SUVR=1.11, 1.21, 1.33, and 1.46) with the AI-imputed tau PET. The lowest and highest cut points (SUVR=1.11 and 1.46) reflecting recent clinical trial stratification^29^ and middle cut points reflecting 95% specificity (SUVR=1.21) and discrimination between age-matched controls and cognitively impaired amyloid PET positive individuals (SUVR=1.33).^50^ In addition, to evaluate the performance of AI-imputed images relative to the input modalities, the ROC analysis was also performed using the actual FDG PET, cortical thickness, or PiB PET as predictors. The cortical thickness was measured with FreeSurfer software.^58^ All variables were derived from the tau PET meta-ROI for the analysis. For FDG PET, we found that applying the model was more successful in predicting tau positivity than the actual FDG SUVR (for FDG, mean AUROC=0.70, 0.78, 0.83, and 0.85 for SUVR thresholds 1.11, 1.21, 1.33, and 1.46, respectively; for FDG-based AI-imputed tau-PET, mean AUROC= 0.70, 0.85, 0.93, and 0.96 for SUVR thresholds 1.11, 1.21, 1.33, and 1.46, respectively; Fig.5A-C). The FDG-based AI-imputed tau PET showed significantly improved AUROC values versus the actual FDG, except for at the lowest SUVR threshold (=1.11) (p=0.004 for 1.21, p<0.001 for 1.33 and 1.46, Holm-Sidak test, Fig.5C). A similar ROC analysis was performed using cortical thickness directly measured from MRI examinations and MRI-based AI-imputed tau PET scans to predict true tau positive participants (Fig.5D-F). As the cortical thickness metric is not combinable across the different manufacturers, the GE and Siemens cohorts were separately analyzed and the result for GE, which is majority manufacturer of our dataset, was displayed in the main result. In contrast to the FDG-based model, the MRI-based AI-imputed tau was not more successful than direct measurement of cortical thickness (for cortical thickness, mean AUROC=0.62, 0.75, 0.82, and 0.88 for SUVR thresholds 1.11, 1.21, 1.33, and 1.46, respectively; for MRI-based AI-imputed tau-PET, mean AUROC= 0.63, 0.78, 0.84, and 0.88 for SUVR thresholds 1.11, 1.21, 1.33, and 1.46, respectively; Fig.5D-F). No significant differences were found in the AUROC (p>0.05 for SUVR thresholds 1.11, 1.21, 1.33, and 1.46, Holm-Sidak test). The Siemens cohort also showed a similar result (Fig.S9). The PiB-based model also showed no significant improvement in the AUROC for tau prediction compared to actual PiB PET SUVR (for PiB, mean AUROC=0.74, 0.83, 0.92, and 0.93 for SUVR thresholds 1.11, 1.21, 1.33, and 1.46, respectively; for PiB-based AI-imputed tau-PET, mean AUROC=0.75, 0.86, 0.94, and 0.96 for SUVR thresholds 1.11, 1.21, 1.33, and 1.46, respectively; Fig.5G-I). This result implies that imputing tau PET scans from MRI and PiB did not add predictive value for classifying tau positivity beyond cortical thickness or PiB PET SUVR alone. In the pairwise comparison between the predictors, FDG- and PiB-based AI-imputed tau PET outperformed the other methods in classifying tau-positivity, except for the lowest cutoff value (Holm-Sidak test, Fig.S10).

**Fig. 5.**
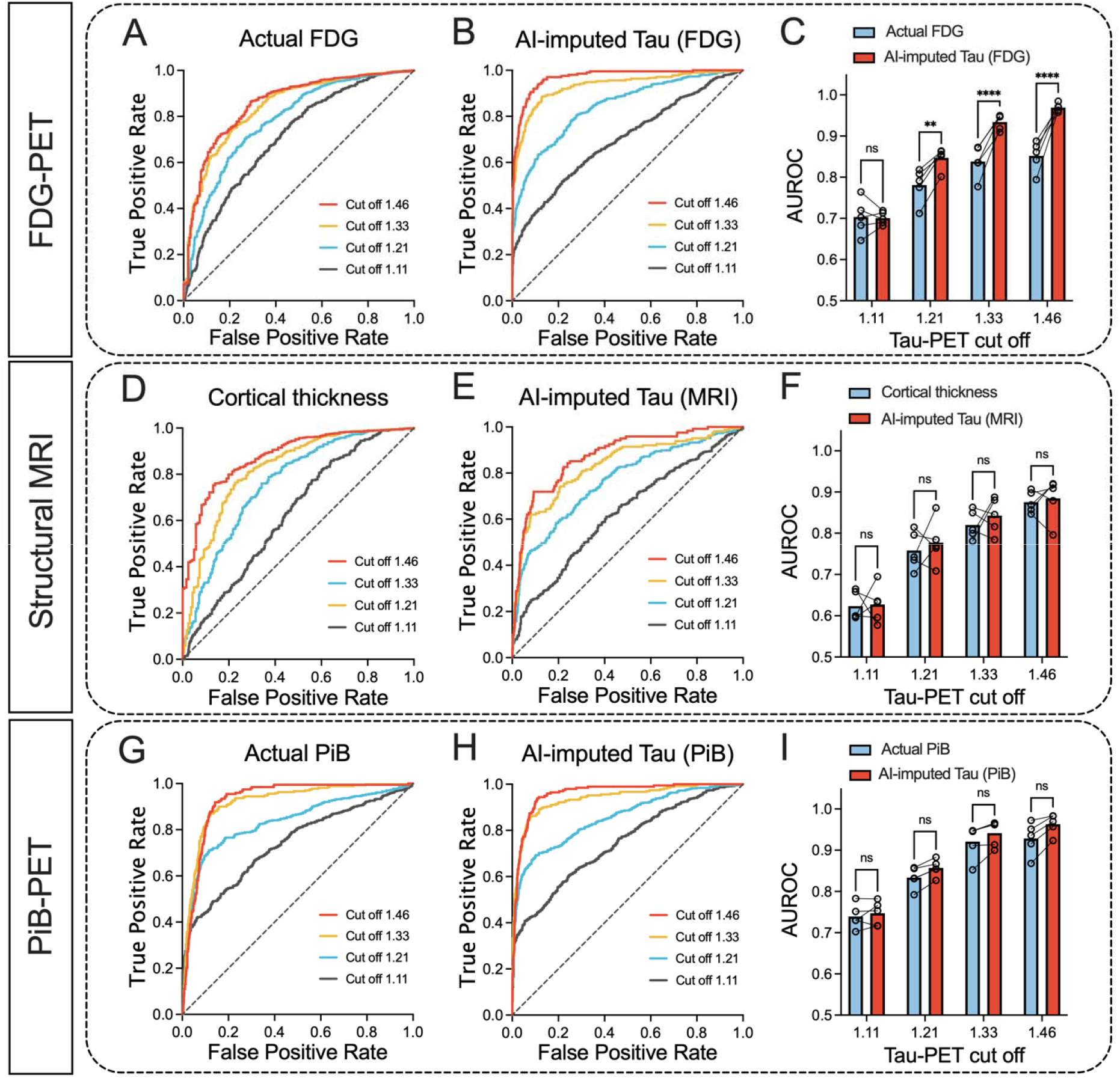
ROC analysis for tau PET positivity. Tau positivity predicted from the groundtruth tau PET using four different meta-ROI cutoff thresholds (1.11, 1.21, 1.33, and 1.46) were obtained using six different predictors: **(A)** Actual FDG PET and **(B)** FDG-based synthesized tau PET with **(C)** AUROC comparison between the original FDG and FDG-based AI-imputed tau PET; **(D)** Cortical thickness from the cohort who had GE scans and **(E)** MRI-based AI-imputed tau PET from the cohort who had GE scans with **(F)** AUROC comparison between the cortical thickness and MRI-based AI-imputed tau PET, and **(G)** Actual PiB PET and **(H)** PiB-based AI-imputed tau PET with **(I)** AUROC comparison between the PiB PET and PiB-based AI-imputed tau PET. A pair-wise comparison was performed between input data and the corresponding AI-imputed tau PET for each cutoff. Statistical significance was tested by post hoc Holm-Sidak comparisons after two-way ANOVA. ** p<0.005, **** p<0.0001. Open circles in C,F and I indicate different folds.

For the ADNI dataset, a similar result was observed (Fig.S11-12). FDG-based AI-imputed tau PET showed significantly improved AUROC values over the actual FDG PET (p<0.001 for 1.11, 1.21, 1.33, and 1.46, Holm-Sidak test, Fig.S11A-C), while the MRI-based model did not show an improved AUROC (p>0.05, for SUVR thresholds 1.11, 1.21, 1.33, and 1.46, Holm-Sidak test, Fig.S12D-F). For all SUVR thresholds, FDG-based AI-imputed tau PET showed the highest AUROC value for classifying tau positivity.

We also trained a CNN model to predict only meta-ROI tau PET SUVR from the cross-modality inputs. For this regression task, a 3D-DenseNet model^59,60^ was utilized with a linear activation and mean absolute error as a loss function. As result, we found that the two different approaches’ performances were not significantly different (Fig.S13A-C). In the ROC analysis for tau positivity, the total volume imputation vs. meta-ROI only imputation did not show a statistically significant difference for every cut off values in all modalities (Fig.S13D-F, p>0.05, Holm-Sidak test).

### Classification Performance of AI-imputed Tau PET for Clinical Diagnostic Group

ROC analysis was also performed to assess the diagnostic performance of AI-imputed tau PET images. For this analysis, four different meta-ROI tau PET values were extracted: actual tau, FDG-based AI-imputed tau, MRI-based AI-imputed tau, and PiB-based AI-imputed tau (Fig.6). For comparison with the model-imputed tau PET, metrics from each input modality were also calculated from tau PET meta-ROI: FDG PET SUVR, cortical thickness, and PiB PET SUVR.

**Fig. 6.**
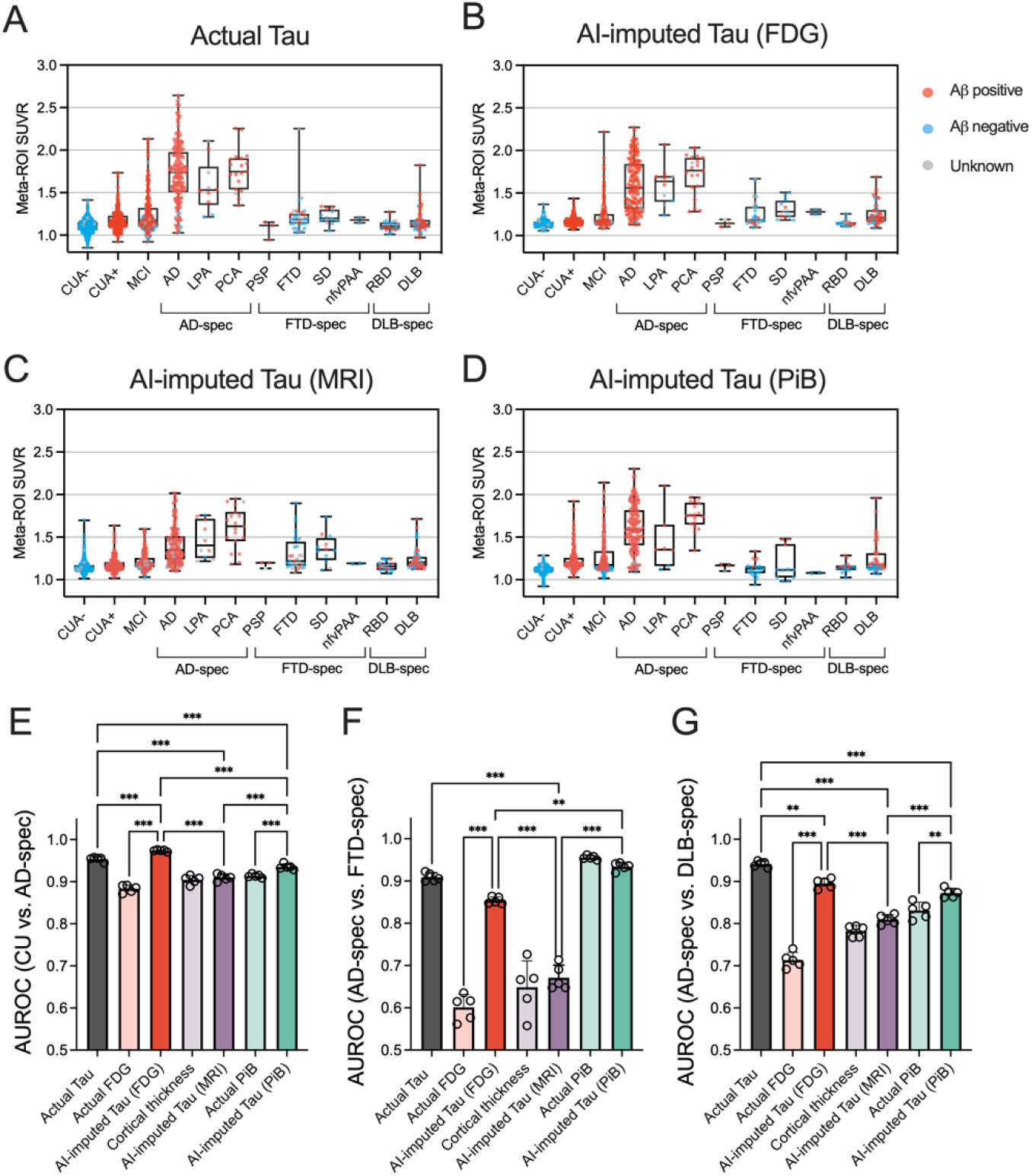
Diagnostic performance of AI-imputed tau PET. **(A-D)** Meta-ROI SUVRs from the actual tau, FDG-, MRI-, and PiB-based AI-imputed tau PET were plotted for each diagnostic group. Red, blue and gray colored dots show amyloid positive, negative, and unknown, respectively. **(E-G)** ROC analysis was performed for classifying the diagnostic groups using seven predictors. Open circles indicate different folds. Statistical significance was assessed with two-way ANOVA and Holm-Sidak post hoc comparison Abbreviations - CUA-: cognitively unimpaired with normal amyloid; CUA+: cognitively unimpaired with abnormal amyloid level; MCI: mild cognitively impaired; AD: Alzheimer’s disease; LPA: logopenic progressive aphasia; PCA: posterior cortical atrophy; PSP: progressive supranuclear palsy; FTD: frontotemporal dementia; SD: semantic dementia; nfvPAA: non-fluent variant of progressive associative agnosia; RBD: REM sleep behavior disorder; DLB: dementia with Lewy bodies. ** p<0.01, *** p<0.0001.

Fig.6A-D show the meta-ROI tau PET SUVR for each diagnostic subgroup. In general, the pattern of distribution was similar across the modalities, while the MRI-based tau PET showed relatively lower predicted SUVR than others (Fig.6C). For the analysis, the diagnostic groups were defined as CU amyloid negative (CUA-) and CU amyloid positive (CUA+), MCI, AD-spec (i.e., AD spectrum including typical AD, LPA, and PCA), FTD-spec (i.e., FTD spectrum including PSP, bvFTD, SD, and nfvPPA), and DLB-spec (i.e., DLB spectrum including RBD and DLB) and the classification was performed for CU vs. AD-spec, AD-spec vs. FTD-spec, and AD-spec vs. DLB-spec (Fig.6E-G). We performed a statistical test for comparisons of AUROC among the AI-imputed and actual tau PET and pair-wise comparison between the AI-imputed tau PET and the metric from the corresponding input data. For the CU vs. AD-spec comparison, the FDG-based model showed the highest accuracy followed by the actual, PiB-based, and MRI-based tau PET (Holm-Sidak test; Fig.6E and H). Interestingly, the classification performance of the FDG-based model was significantly higher than the actual tau PET (p<0.001, Holm-Sidak test, Fig.6E). In comparison with input modality, FDG- and PiB-based model showed improved accuracy (p<0.001, Holm-Sidak test, Fig.6E). For the AD-spec vs. FTD-spec, the PiB-based model showed the best performance followed by the actual, FDG-based, and MRI-based tau PET (Fig.6F); however, the PiB PET also performed well and was not significantly different with the synthesized tau PET (p=0.65, Holm-Sidak test, Fig.6F). The performance of FDG-based tau model was significantly improved compared to the FDG PET (p<0.001, Holm-Sidak test, Fig.6F). In classifying the AD-spec vs. DLB-spec, the actual tau performed the best, followed by the FDG-based, PiB-based, and MRI-based AI-imputed tau PET (Fig.6G). FDG-and PiB-based model showed an improvement upon the performance of the input data (p<0.001 and p=0.007 for FDG-based model and PiB-based model, respectively, Holm-Sidak test, Fig.6G).

### Interpretability of 3D dense-U-net Model using Occlusion Sensitivity Analysis

To facilitate the interpretability of the dense-U-net model, saliency maps were estimated through occlusion sensitivity analysis for three different input modalities.^60,61^ In the occlusion method, a single ROI in the input space was occluded by setting these voxels to zero, and their relevance in the decisions was indirectly estimated by calculating the change of MAPE (*i.e.*, ΔMAPE = MAPE_occlusion_ - MAPE_original_). An adjacency matrix (Fig.7A, C, and E) plotting the regional ΔMAPE against each occluded ROI (vertical axis) shows the contribution of each ROI to the performance of the model. The diagonal line in each of these adjacency matrices is somewhat expected, representing the high contribution of the voxel of source images to the same region of the synthesized tau, observed for all three modalities (Fig.7A,C, and E). Notably, occlusion analysis revealed multiple additional anatomic regions with a high contribution that are spatially remote which differ according to the input images. For the FDG-based model, the sensorimotor cortex and the frontal lobe were dominant contributors to the global accuracy of the tau model, showing a high contribution to the MAPE for most of the brain (Fig.7A and B). This implies that metabolism in the sensorimotor cortex and the frontal regions were involved in the accurate imputation of tau PET for other brain regions. On the other hand, for the MRI-based model, the temporal, parietal, and occipital lobes were found to be the dominant contributor to global accuracy (Fig.7C and D). The influence of remote structures was less prominently observed in the PiB-based model (Fig.7E and F), implying that this model generates the AI-imputed tau PET images using only relatively local amyloid information. For every model tested, no interhemispheric effect was convincingly observed.

**Fig. 7.**
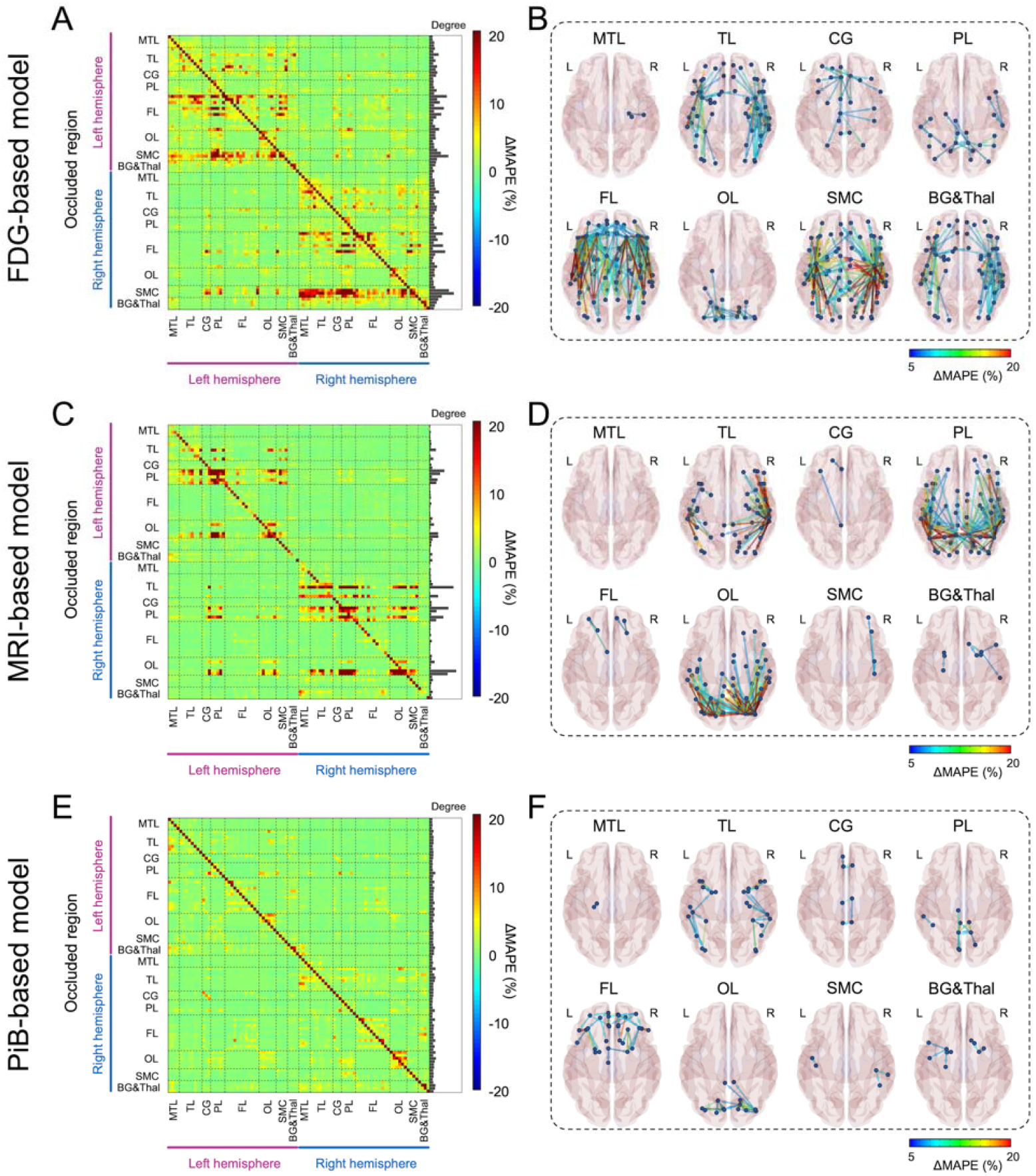
Occlusion analysis. ROI-wise occlusion analysis was performed to enhance the interpretability of model. **(A, C, and E)** The adjacency matrix shows the ΔMAPE in one ROI (horizontal axis) from occluding another ROI (vertical axis) for FDG-, MRI-, and PiB-based model, respectively. ΔMAPE was calculated as MAPE_R1→R2_ - MAPE_R2_, where R1 is an occluded ROI and R2 is the region where the MAPE is calculated. The right panel in each matrix indicates the summation of ΔMAPE along the horizontal axis. **(B, D, and F)** 3D rendering plots of the adjacency matrix in A, C, and E for FDG-, MRI-, and PiB-based model, respectively. Each edge’s color was illustrated by ΔMAPE value between nodes. Each label denoted above the figure indicate the occluded regions. Abbreviations - MTL: medial temporal lobe; TL: temporal lobe; CG: cingulate cortex; PL: parietal lobe; FL: frontal lobe; OL: occipital lobe; SMC: sensorimotor cortex; BG&Thal: basal ganglia and thalamus

## DISCUSSION

We showed that 3D dense-U-net models can successfully impute tau PET using cross-modality imaging input. Overall, FDG-based approaches showed the highest degree of accuracy with good correlation to ground-truth tau PET and low error for regional SUVRs, followed by the PiB-based model. The MRI-based model was marginally accurate, but significantly inferior to the FDG- and PiB-based models. In addition, the FDG-based model showed the most robust prediction capability, performing accurately in an external cohort from the ADNI database where the MRI-based model did not. In testing the clinically relevant application of AI-imputed tau PET to predict tau positivity and classify diagnostic groups, only the FDG-based model showed significant improvement upon the performance of the original input data, suggesting that the model may enhance the utility of the metabolic images alone. The occlusion method, employed in an attempt to allow interpretation of the model’s mechanism of prediction, revealed that the FDG- and MRI-based models utilized global input from physically remote ROIs to impute the tau PET, whereas a relatively locoregional contribution was predominantly observed in the PiB-model.

We speculate that the dense-U-Net models generated tau PET images using the patterns of hypometabolism, cerebral atrophy, and amyloid burden captured by FDG PET, structural MRI, and PiB PET, respectively. The possibility that FDG hypometabolism, atrophy, and amyloid levels are important features of the model, facilitating the successful imputation of tau PET images, is biologically plausible and supported by previous literature. A strong correlation of the tau uptake on tau PET and hypometabolism on FDG PET is well documented in prior studies.^11,19,33,62^ The regional atrophy pattern identified on MRI correlates well with regional tau PET uptake.^13,20,34,35,63^ Autopsy studies also support a strong correlation of tau burden and brain atrophy.^64–67^ A correlation between tau and amyloid distribution has been shown, although the molecular relationship is complex, with a stronger relationship observed in the temporoparietal regions to a greater degree for predominantly cognitively normal cohorts^34,36–38^ and in the frontal, parietal, and occipital lobes in more advanced dementia cohort.^25^ These neuroimaging AD biomarkers become abnormal in a temporally ordered manner.^68^ The amyloid PET tracer uptake increases earliest followed by tau PET and FDG PET, then structural MRI, and finally clinical symptoms. The amyloid cascade hypothesis suggested that accumulation of Aβ plaques is the primary cause of tau NFT formation^69^; however, it also has been suggested that the aggregation of toxic form of Aβ and tau might be independent processes separately contributing to the development of AD pathology.^70^ In addition, autopsy data has shown that the regional patterns of Aβ differ from that of tau deposition.^56^ Meanwhile, hypometabolism and atrophy are more closely related to tau accumulation as a downstream consequence of neuronal loss due to tau NFTs. Abnormalities on FDG-PET may occur before structural changes in the brain in AD^71,72^ and hence potentially closer in time to the tau deposition, perhaps relating to the better performance of the FDG-based imputation in our study. Whitwell et al. also showed that FDG hypometabolism correlated with tau PET uptake better than cortical thickness or PiB in both typical AD and atypical AD, implying that FDG metabolism is most sensitive to the effects of tau pathology.^62^ This is concordant with our observation that the FDG-based model was more successful than the MRI- and PiB-based model in comparison to ground-truth tau PET. FDG has also been proposed as a marker of other conditions of interest in clinical dementia populations beyond those associated with AD-tau, such as hippocampal sclerosis and TDP-43,^73^ DLB,^74–76^ and is currently approved and covered by Medicare for clinical use in the differential diagnosis of FTD and AD.^77^ This wide applicability across the spectrum of clinical conditions included in building our model is likely a key contributing factor in our models potential to enhance the clinical utility of FDG.

Furthermore, occlusion analysis revealed that the model’s predictions of tau activity for a given ROI depends not only on that same anatomic ROI but also on global input from physically remote ROIs. This held true for both the FDG- and MRI-based model’s predictions, whereas the PiB-based model demonstrated predominantly locoregional contributions to the tau prediction for each ROI. The result implies that local amyloid levels alone may be sufficient for the PiB-model to predict accurate synthesized tau images while the trained model did not simply impute tau PET using local FDG PET activity or MRI features but relied on additional associations to inform the distribution of tau in the model-imputed PET images. This also provides clues about the enhanced performance of FDG-based model compared to the actual FDG in contrast to the other models did not.

For the FDG-based model, one possible interpretation is that the model could predict the tau level based on SUVR comparisons between ROIs. The primary sensorimotor cortex and frontal lobe were dominant areas of influence on synthesized tau PET accuracy from the FDG-based model revealed by occlusion analysis, surprising because the sensorimotor cortex is typically spared from FDG hypometabolism and frontal lobe involved in later stages of Alzheimer’s dementia.^78^ Therefore, preserved metabolism in these regions may be interpreted by the model as reference region, modifying the model’s prediction of tau uptake in remote locations of the brain. In this context, it is important to note that sparing of the sensorimotor strip in AD is a feature currently used by clinicians to inform expert visual interpretation of FDG PET images.^79^ This clinically used feature is thought to be related to AD biology rather than image intensity normalization. This feature, juxtaposed with heteromodal association cortex, also characterizes principal patterns of functional connectivity^80^ that are also observed in modes of variation in FDG PET related to global functional architecture across dementia syndromes.^31^ Therefore, one biologically plausible alternative explanation is that the model could utilize information related to the brain’s global functional architecture in predicting the local tau uptake. Previous works have supported the relationship between tau pathology and brain connectivity, based on study of tau PET distribution and correlation to resting-state functional MRI (fMRI).^12,21,81–83^ Franzmeier et al., also reported that the higher functional connectivity observed by the resting-state fMRI is associated with higher rates of tau accumulation.^84^ However, different functional properties have been associated with amyloid^83^ which may potentially explain the difference in learned features for these two modalities in predicting tau pathology in our study. This is consistent with our recent study showing that the FDG-PET based global functional state space showed a much higher predictive accuracy for tau PET and Braak NFT stage than amyloid PET.^31^ A key feature of the global functional state space described in that study, is the juxtaposition of heteromodal association cortex with primary sensorimotor cortex, and this feature is reflected in FDG occlusion analysis (Fig.7A-B), but not in the amyloid occlusion analysis (Fig.7E-F) observed in the current study.

For the MRI-based model, the dominant regions of influence on remote parts of the brain were in the temporal, parietal, and occipital lobes, correlating to areas of characteristic tau deposition and areas of significant regional volume loss in the AD spectrum.^7^ Similar to the FDG-based model, the MRI model utilized the global information to predict the tau level; however, applying the model did not improve the performance compared to cortical thickness. The marginal predictive accuracy of the MRI model, presumably hampered by the heterogeneity of structural changes, may not be robust enough to improve upon the structural images which have higher spatial resolution compared to the PET scans. Structural imaging is also less sensitive to changes in functional networks than FDG PET and therefore potentially less likely to contain the same level of functional network information on a single subject level that could be used to predict tau PET.

Off-target binding of the tau tracer AV-1451 is an incompletely characterized phenomenon, most frequently described in the substantia nigra, caudate, putamen, and choroid plexus on the basis of post-mortem analysis and autoradiography studies.^15,85,86^ The literature on the topic has suggested non-specific binding to structurally similar molecules such as MAO-A, MAO-B and potentially to mineralized or pigment-containing structures such as neuromelanin. Because the off-target binding of AV-1451 is not correlated to hypometabolism, atrophy, and amyloid-beta burden, we would expect the off-target binding to be somewhat poorly predicted in the synthetic tau images and this is what we observe. The common locations of off-target binding that were included in the ROI analysis demonstrate relatively low correlation and high MAPE with the imputed-tau PET scans and ground truth for all of the models (Fig.2–4). The basal ganglia regions showed some association between ground-truth and AI-imputed SUVR (Fig.S14). Previous studies have shown that the non-specific binding in the basal ganglia is associated with age, as the neuromelanin and iron increases with age.^15,87^ Thus, we speculate that the AI model may impute the off-target binding by learning agerelated changes of input.

The AI-imputed tau PET is also limited, to a certain extent, by the properties of true ^18^F-Flortaucipir PET. The model follows the behavior of AV-1451 PET, used as the ground-truth, and not necessarily the distribution of tau that might be found at autopsy. This is also a strength, in that the model generates a result that is analogues to a clinically useful diagnostic procedure; however, this also has some complex implications. The AV-1451 tracer varies in strength of binding to tau isoforms, binding less to 3R or 4R tau than 3R+4R tau.^15^ This is also reflected in vivo suggesting more specificity of AV-1451 for AD-like tau than other tau isoforms.^28^ Numerous studies have confirmed the role of AV-1451 in detecting non-Alzheimer’s tauopathies is limited.^88,89^ Interestingly, the FDG-based model’s overall prediction error was slightly higher in the temporal region for the FTD cohort and parietal region for the DLB cohort than the AD cohort (Fig.S15), which are the regions with characteristic hypometabolism in each disease.^79,90^ This may in part reflect a less direct relationship between areas impacted by these isoforms, changes of FDG PET, and the predicted tau PET activity. Nonetheless, the diagnostic performance of the model’s meta-ROI for these groups of FTD and DLB participants was generally accurate and significantly enhanced the performance of FDG PET alone. The differences between the input and output of the model and the regional variation with different types of tau-pathology support our speculation that the model is not directly “translating” FDG uptake into tau for specific region but is more likely utilizing global input from physically remote ROIs and broader pattern-recognition mechanisms to predict tau activity for a given region.

The AI-imputed tau PET may allow clinicians and researchers to maximize the use of neuroimaging biomarkers with a projection of tau pathology. Particularly, application of the model would be most beneficial for FDG PET to enhance the utility of the metabolic images that are being obtained in the evaluation of these patients currently. The high correlation to ground-truth, including in an external dataset, implies that AI-imputed tau PET may be a viable alternative of tau PET in scenarios where tau PET is not feasible, or the tau radiotracer is unavailable. As outlined in the introduction, use of multiple radiopharmaceuticals for FDG, tau, and amyloid PET is an extremely expensive and resource-intensive prospect, now emerging as an area of research and clinical need with recent FDA accelerated approval of Alzheimer’s disease-modifying therapy^91^ and the potential for additional targeted therapies in the future. In contrast to tau PET agents, [^18^F]FDG PET is one of the most widely-utilized imaging biomarkers for AD,^92^ is clinically used to characterize multiple types of dementia.^93^ FDG PET has the support of multiple professional societies in the diagnosis of dementia and is accessible at many medical centers ^94–100^ Our study demonstrates that FDG-imputed tau PET may provide valuable information regarding tau pathology with a high correlation to real tau PET, which could have applications in settings where only FDG PET is feasible, and potential for use as a biomarker in the research setting. Ultimately, AI-imputed tau PET may enable more efficient resource utilization and reduce patients’ exposure to multiple radiopharmaceuticals and imaging tests by maximizing the information gleaned from FDG PET.

One important limitation of the study is that the PiB-based model was not cross-evaluated in the external dataset, as amyloid PET using the PiB radiotracer was not available from the ADNI database. While the CNN model was tested in participants from three different cohorts, and externally validated in the ADNI dataset for the FDG PET and MRI-based input, it remains unknown how the model will perform using input from more diverse imaging settings and patient populations. While the occlusion sensitivity analysis provides some insight about which regions are important to the success of the model, the mechanism of projecting tau uptake by the model remains unknown, hindering our ability to make inferences about the relationships between brain structural changes, tau and amyloid deposition, and glucose metabolism. We use the information here to imply relationships between tau deposition and metabolism in other parts of the brain which merit exploration with further research. We acknowledge that these measurements include small regions, where measurements from PET images are nosier than in larger regions, but this limitation is common to most published analyses of tau PET.

In summary, a 3D Dense-U-Net architecture is presented which produced synthesized tau PET brain scans from FDG PET, PiB PET, and MRI. The FDG-based model of AI-imputed tau PET demonstrated a high degree of correlation to ground-truth tau PET for patients on the MCI-AD spectrum, distinguished tau-positive versus tau-negative patients, and classified diagnostic groups with performance similar to the AV-1451 tau PET exams. AI-imputed tau is feasible and has a potential to augment the value of FDG PET for patient MCI and AD patients. Additional work to utilize multi-modality imaging input simultaneously is also needed, as this may further enhance the accuracy of model-imputed tau PET.

## MATERIALS AND METHODS

### Participants

Participants from the Mayo Clinic Study of Aging (MCSA) or the Alzheimer’s Disease Research Center (ADRC) study who underwent MRI, FDG PET and Tau PET were included (n=1,192, number of scans=1,505)(Table 1 and Fig.S1). All participants or designees provided written consent with the approval of Mayo Clinic and Olmsted Medical Center Institutional Review Boards. The participants were categorized into major clinical sub-groups based on clinical diagnosis including CU (n=739, number of scans=890), MCI (n=169, number of scans=208), typical AD (n=110, number of scans=165), bvFTD (n=25 number of scans=32), DLB (n=38, number of scans=54) and other clinical syndromes (e.g., vascular cognitive impairment, RBD, PCA, SD, lvPPA, nfPPA and PSP; n=111, number of scans=156)(Table 1). To examine whether the trained model presents a dataset-specific bias, we also utilized the Alzheimer’s Disease Neuroimaging initiative (ADNI; adni.loni.usc.edu) dataset. For the ADNI cohort, we pulled all visits with a 3T accelerated T1-MRI, FDG PET and Tau PET where available (n=288; table S3). The ADNI dataset included normal controls (n=15), MCI (n=205) and Dementia (n=68). Image IDs for the ADNI cohort used in this study can be downloaded from the following link (https://github.com/Neurology-AI-Program/AI_imputed_tau_PET/ADNI_cohort_with_imageIDs.csv).

### Neuroimaging

For the Mayo data, T1-weighted MRI scans were acquired using 3T GE and Siemens scanners with MPRAGE sequences. FDG PET imaging was performed with ^18^F-FDG, amyloid PET with ^11^C-PiB^44^ and tau PET with ^18^F-Flortaucipir (AV-1451).^45^ PET images were acquired 30-40 minutes after injection of F-18 FDG, 40-60 minutes after PiB injection and 80-100 minutes after AV14151 injection. CT was obtained for attenuation correction. Details of ADNI imaging protocols have been previously published.^101,102^ PET images were analyzed with our in-house fully automated image processing pipeline.^103^ Briefly, the PET scans were co-registered to the corresponding MRI for each participant within each timepoint, and subsequently warped to Mayo Clinic Adult Lifespan Template (MCALT) space which has dimensions 121×145×121 voxels^48^ (https://www.nitrc.org/projects/mcalt/) using the nonlinear registration parameters from SPM12 unified segmentation. The corresponding MRI was corrected for intensity inhomogeneity and segmented using MCALT tissue priors and segmentation parameters. FDG PET standardized uptake value ratio (SUVR) was calculated by dividing the median uptake in the pons and the SUVR images were used for input data to the CNN model. Tau PET SUVR was calculated by dividing the median uptake in the cerebellar crus grey matter^104^ and the tau PET SUVR images were used for target label data for CNN training. For each MRI volume, an intensity normalization was performed by dividing a mean intensity derived from individualized white matter mask.^105^ Cortical thickness was measured with FreeSurfer software, version 5.3.^58^ The tau PET meta-ROI used in this study included the amygdala, entorhinal cortex, fusiform, parahippocampal and inferior temporal and middle temporal gyri.^104,106^ The meta-ROI SUVR was calculated as an average of the median uptake across regions of meta-ROI.

### Network Architecture

A schematic of the 3D Dense-U-Net architecture used for this study is shown in Fig. 1.^46^ The network is a U-Net type architecture^47^ with dense interconnections between convolutional layers (dense block). The architecture is comprised of 4 down-sampling (encoder) blocks for feature extraction and 4 up-sampling (decoder) blocks for image reconstruction, which are connected by a bridge block. Each encoder block has two padded convolutions (3×3×3) with a rectified linear unit (ReLU) as an activation function, followed by a 2×2×2 max pooling with stride 2 for down-sampling. In the decoding path, each block is comprised of a 2×2×2 up-convolution with stride 2 for up-sampling,^107^ followed by two padded convolutions (3×3×3) with a ReLU activation function. The output of the last convolutional layer, prior to the pooling operation of each encoder block, is concatenated with the output of the up-pooling layer in the associated decoder block through a skip connection. The bridge block (in the lower part of the network) is composed of two padded convolutions (3×3×3) with a ReLU activation function. Within every block, the convolutional layers are densely interconnected in a feed-forward manner (illustrated in the left dotted box in Fig. 1). The network doubles or halves the number of filters (denoted above each block) along each successive encoder and decoder path, respectively. After the last up-sampling dense block, a single 3×3×3 convolutional layer with linear activation is used to generate an output image.

The architecture takes input volumes of size 128×128×128 and outputs the images with the same dimensions. For this, we resized the volume by cropping and zero padding so that it is divisible by 2 until the bottom of the network for the max-pooling and upsampling procedure. Along the AP axis, 8 slices of anterior and 9 slices of posterior were cropped. Then, 7 slices were padded on the left and bottom of the cropped volume, forming a 3D data of size 128×128×128. The output volume of the network was reconstructed as the original size (121×145×121) for the visualization.

### Training and Testing

The neural network was implemented using Keras with Tensorflow as the backend. Cross-validation experiments were conducted using 5-fold validations (60% training set, 20% validation set, and 20% test set). In order to prevent any possible data leakage between the training and validation/testing datasets, we excluded any overlap of participants among training, validation, and test sets. Within each set, multiple scans per subject were included. During training, the model was optimized using Adam optimizer^108^ with parameters: β1=0:9 and β2=0.999. The training epoch used was 150. The learning rate from training was set to 1×10^-4^ and decreased by a factor of 2 for every ten epochs. If the validation error did not improve in 7 epochs, the learning rate was updated. We had used a mini-batch of size 2. The mean squared error was used as the loss function. The training and testing are performed on Tesla P100 GPU. The source code are available online: https://github.com/Neurology-AI-Program/AI_imputed_tau_PET.

### Occlusion analysis

Occlusion sensitivity analysis was performed to identify regions of interest in the brain contributing to the performance of the tau PET synthesis model.^61^ The analysis was conducted for the test image data set for FDG-, MRI- and PiB-based models. For each model, voxels from a single ROI in the original source images were occluded with zero values one at a time, and their relevance in the tau PET synthesis was estimated as a change of regional MAPE between the original and after occlusion (ΔMAPE_R1→R2_= MAPE_R→R2_ - MAPE_R2_, where R1 is an occluded ROI and R2 is a region where the MAPE is calculated).

### Neuropathology methods

The neuropathologic assessment was performed as reported previously.^8^ Briefly, brain sampling and standardized neuropathologic examination were performed according to the CERAD protocol.^109^ Tissue samples were paraffin-embedded and 5 μm thick tissue sections were routinely stained with hematoxylin and eosin, as well as a modified Bielschowsky silver stain. Immunohistochemistry was performed using a phosphospecific tau antibody (AT8; 1:1000; Endogen, Woburn, MA). The AT8 immunostained sections were used to assess Braak tangle stage and neuritic plaque score.^54,55^ Participants were assigned the neuropathologic diagnosis of AD if they had a Braak tangle stage of ≥ IV and had at least a moderate neuritic plaque score. Primary agerelated tauopathy (PART) was assigned if the case met published criteria – Braak tangle stage I-IV and Thal amyloid phase 2 or less. Lewy body disorders were classified neuropathologically based on the distribution and severity of Lewy bodies and neurites^72^.

### Statistical analysis

To evaluate the model’s performance, regional SUVRs were extracted from both the ground-truth and AI-imputed tau PET scans and the Pearson’s correlation and mean absolute percentage error (MAPE) between tau images across participants were tested. A difference of correlation coefficient and MAPE between the models was evaluated using a two-way ANOVA. Receiver operator characteristic (ROC) analyses were performed to compare the discriminatory power of AI-imputed tau PET for tau positivity. The tau positivity was defined using four different meta-ROI cutoff thresholds (1.11, 1.21, 1.33, and 1.46).^29,50^ Six different predictors were utilized: original FDG PET, FDG-based AI-imputed tau PET, cortical thickness, MRI-based AI-imputed tau PET, original PiB PET and PiB-based AI-imputed tau PET. Within the modality type, a pair-wise comparison of the area under the ROC (AUROC) value was performed using two-way ANOVA and Hold-Sidak *post hoc* test. These analyses were performed using the entire cohort; however, the MRI-related variables (i.e., cortical thickness and MRI-based AI-imputed tau) were separately analyzed by their manufacturer (GE and Siemens) because combining the cortical thickness values across the manufacturer is not reliable. For each cutoff value, a pair-wise comparison of the AUROC was performed using a one-way ANOVA with Holm-Sidak *post hoc* test. The ROC analyses were also performed to evaluate the classification performance of tau PET images for multiple diagnostic groups. The diagnostic groups were defined as CU (CUA- and CUA+), AD-spec (AD, LPA, PCA), FTD-spec (PSP, FTD, SD, nfvPPA), and DLB-spec (RBD and DLB) and the analysis was performed for CU vs. AD-spec, AD-spec vs. FTD-spec, and AD-spec vs. DLB-spec. A pair-wise comparison of the AUROC value was performed using a two-way ANOVA and Holm-Sidak *post hoc* test. To document the transparency of the trained model, we developed a model card accompanying benchmarked evaluation in a variety of conditions, such as across different race, ethnicity, and demographics. The model card can be downloaded from the following link (https://github.com/Neurology-AI-Program/AI_imputed_tau_PET.git).

## Acknowledgments

We gratefully acknowledge the support of NVIDIA Corporation with the donation of the Tesla P100 GPU used for this research. We would also like to acknowledge the help provided by the Google team (Stephen Pfohl, Gerardo Flores, Karan Shukla, Katherine Heller, and Parker Barnes) for developing the model card.

## Author contributions

Conceptualization: JL, HM, DTJ

Methodology: JL, MLS, CTM, HJW, ESL, LRB, JLG, CGS, DTJ

Investigation: JL, BJB, HM, HB, JG, LRB, JLG, CGS, KK, DSK, BFB, VJL, RCP, CRJ, DTJ

Visualization: JL

Funding acquisition: CRJ, RCP, DTJ

Supervision: HM, HB, VJL, CRJ, DTJ

Writing – original draft: JL, BJB

Writing – review & editing: All authors.

All authors have given final approval of this version of the article.

## Data and materials availability

The data that support the findings of this study are not publicly available. Data may be available from the authors upon reasonable request and with permission. The source code is available online (https://github.com/Neurology-AI-Program/AI_imputed_tau_PET).

## Supplementary Materials

Fig. S1. Data inclusion/exclusion criteria.

Fig. S2. Comparison of histogram for meta-ROI SUVR among train, validation and test set.

Fig. S3. Testing the FDG-based model on the ADNI dataset.

Fig. S4. Testing the MRI-based model on the ADNI dataset.

Fig. S5. Voxel-wise error maps and multi-scale structural similarity index.

Fig. S6. Comparisons of performance among the AI-imputed tau PET and ground truth tau PET in SliceView.

Fig. S7. 3-dimensional stereotactic surface projection images.

Fig. S8. AI-imputed tau PET and Braak stage.

Fig. S9. ROC analysis for the Siemens cohort.

Fig. S10. AUROC comparisons for tau positivity.

Fig. S11. ROC curves showing the classification performance on the tau positivity tested on ADNI dataset.

Fig. S12. AUROC comparisons for different MRI manufacturers tested on ADNI dataset.

Fig. S13. Comparison between the total volume imputation vs. meta-ROI only imputation.

Fig. S14. AI-imputed SUVR of regions of off-target bindings.

Fig. S15. Regional MAPE distribution for FDG-based AI-imputed tau-PET for AD, FTD, and DLB diagnostic groups.

Table S1. Demographics for FDG/MRI model

Table S2. Demographics for PiB model

Table S3. ADNI Cohort Demographi

