## Supplementary Materials for "Synthesizing Images of Tau Pathology from Cross-modal Neuroimaging using Deep Learning"

Fig. S1. Data inclusion/exclusion criteria.

Fig. S2. Comparison of histogram for meta-ROI SUVR among train, validation and test set.

Fig. S3. Testing the FDG-based model on the ADNI dataset.

Fig. S4. Testing the MRI-based model on the ADNI dataset.

Fig. S5. Voxel-wise error maps and multi-scale structural similarity index.

Fig. S6. Comparisons of performance among the AI-imputed tau PET and ground truth tau PET in SliceView.

Fig. S7. 3-dimensional stereotactic surface projection images.

Fig. S8. AI-imputed tau PET and Braak stage.

Fig. S9. ROC analysis for the Siemens cohort.

Fig. S10. AUROC comparisons for tau positivity.

Fig. S11. ROC curves showing the classification performance on the tau positivity tested on ADNI dataset.

Fig. S12. AUROC comparisons for different MRI manufacturers tested on ADNI dataset.

Fig. S13. Comparison between the total volume imputation vs. meta-ROI only imputation.

Fig. S14. AI-imputed SUVR of regions of off-target bindings.

Fig. S15. Regional MAPE distribution for FDG-based AI-imputed tau-PET for AD, FTD, and DLB diagnostic groups.

Table S1. Demographics for FDG/MRI model

Table S2. Demographics for PiB model

Table S3. ADNI Cohort Demographics

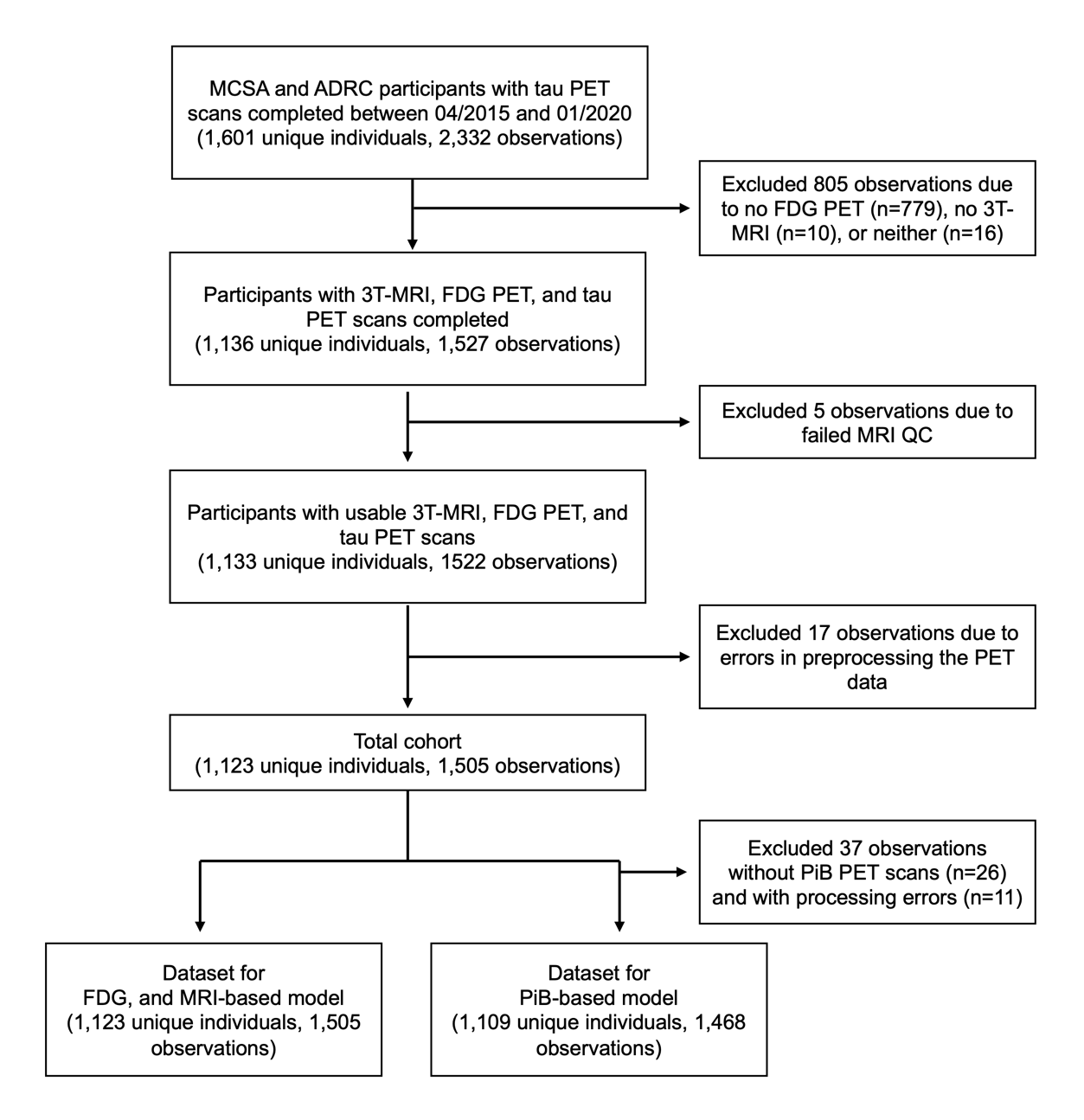

**Supplementary figure 1. Data inclusion/exclusion criteria.**

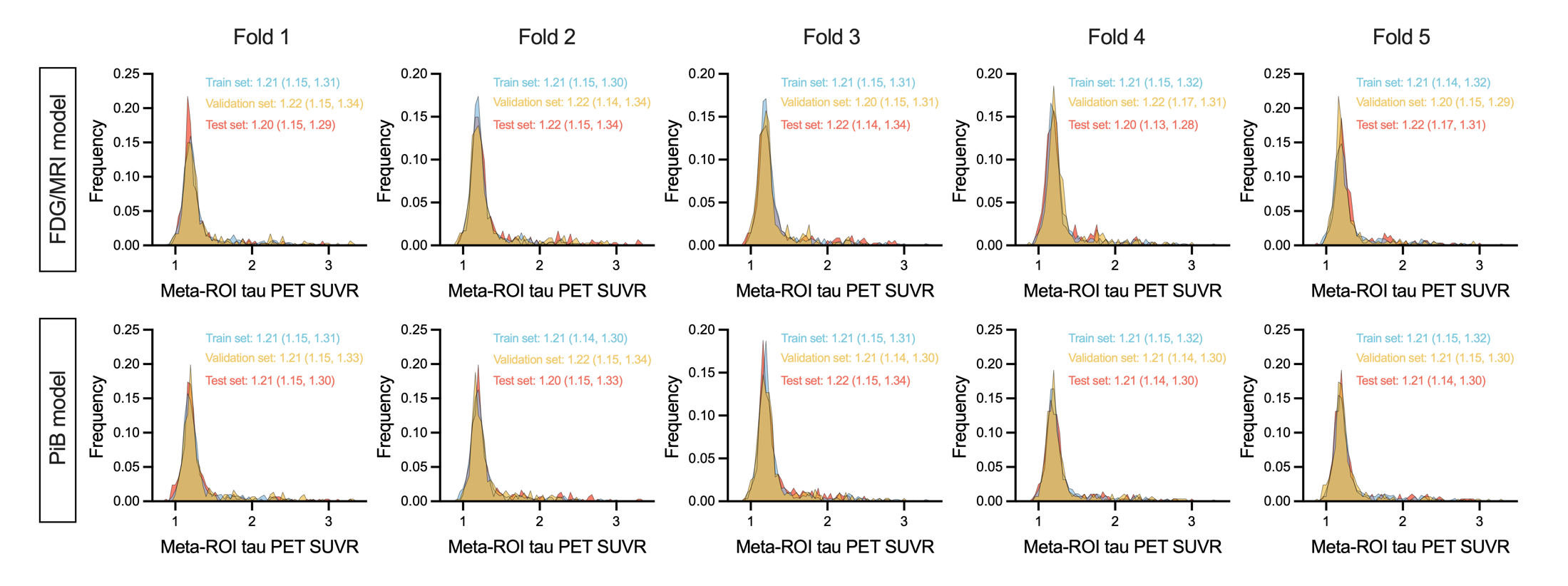

**Supplementary figure 2. Comparison of histogram for meta-ROI SUVR among train, validation and test set.** For each histogram, the values represent median (Q1 and Q3).

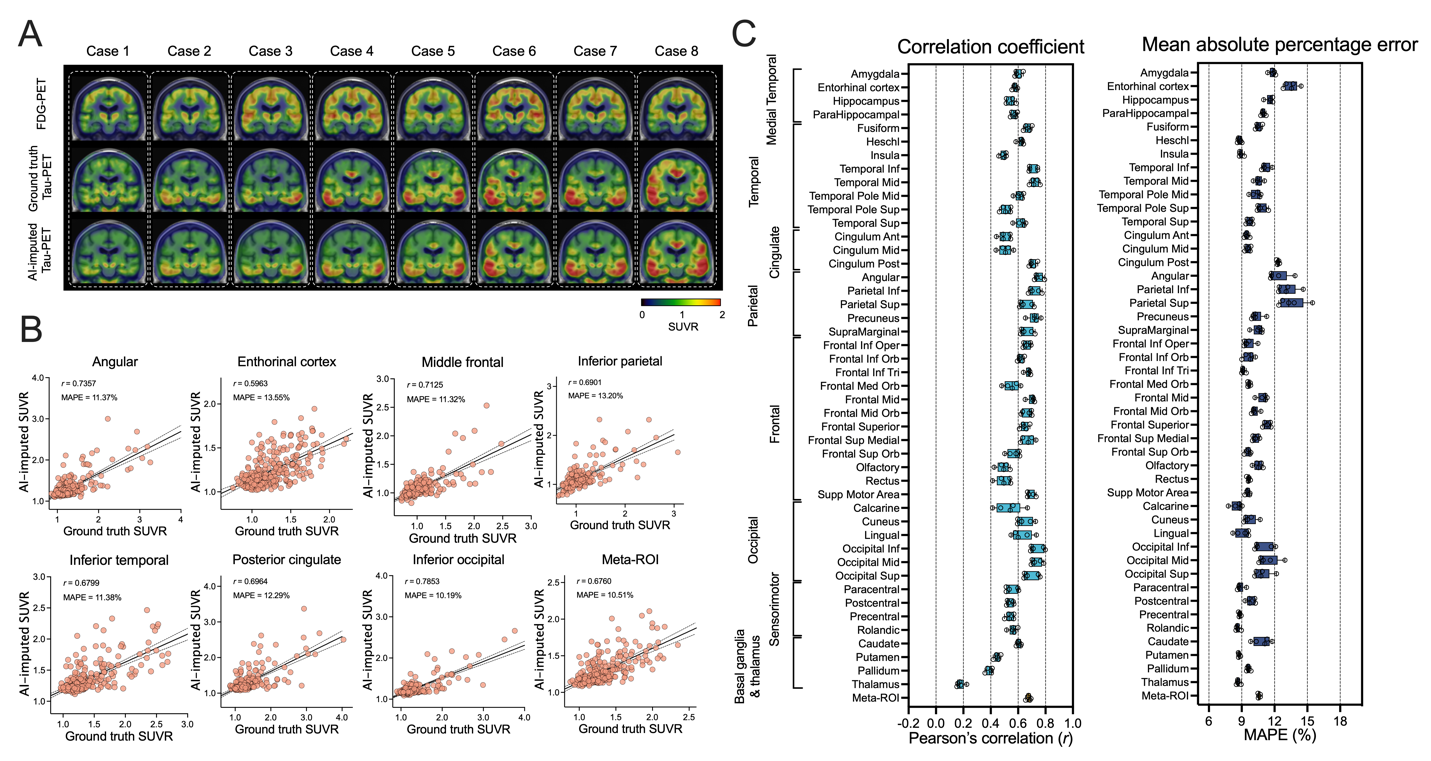

**Supplementary figure 3. Testing the FDG-based model on the ADNI dataset.** (A) Eight representative cases with original FDG-PET, ground truth tau PET and AI-imputed tau PET. (B) Scatter plots of ground truth tau PET and AI-imputed tau PET from seven representative ROIs and meta-ROI. *r* indicates the Pearson’s correlation coefficient and MAPE indicates mean absolute percentage error. Linear regression (black line) and 95% confidence bands (dotted lines) are shown. (C) The correlation coefficient and MAPE from 46 ROIs and meta-ROI is summarized in a box plot. The yellow-colored box depicts the meta-ROI result.

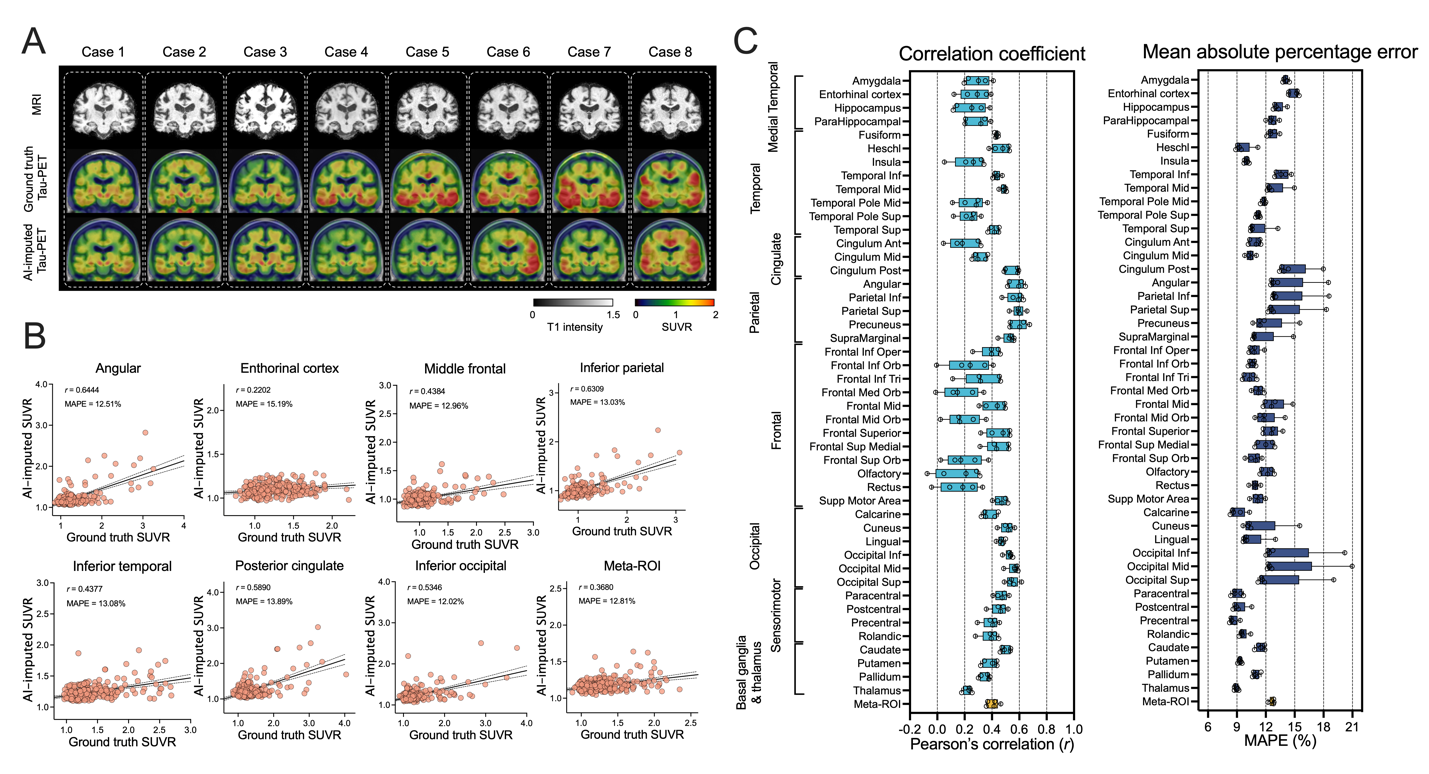

**Supplementary figure 4. Testing the MRI-based model on the ADNI dataset.** (A) Eight representative cases with original MRI, ground truth tau PET and AI-imputed tau PET. (B) Scatter plots of ground truth tau PET and AI-imputed tau PET from seven representative ROIs and meta-ROI. *r* indicates the Pearson’s correlation coefficient and MAPE indicates mean absolute percentage error. Linear regression (black line) and 95% confidence bands (dotted lines) are shown. (C) The correlation coefficient and MAPE from 46 ROIs and meta-ROI is summarized in a box plot. The yellow-colored box depicts the meta-ROI result.

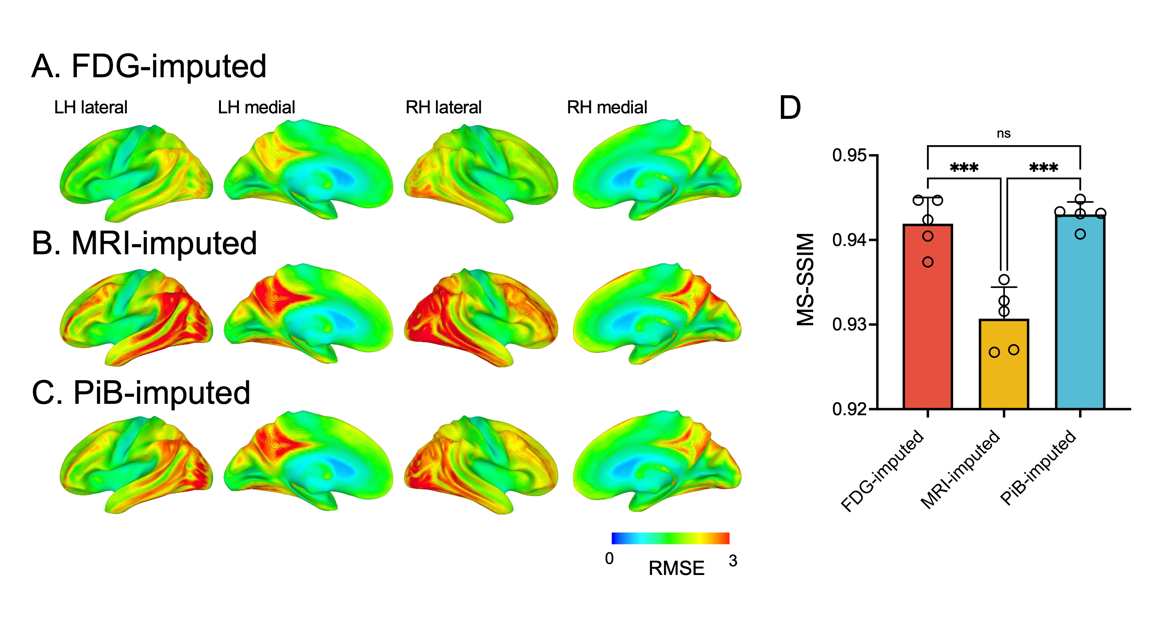

**Supplementary figure 5. Additional evaluation of the model’s performance.** (A-C) 3D rendered images of the voxel-wise root mean squared error (RMSE) map. (D) multi-scale structural similarity index for FDG-, MRI-, and PiB-based models. Statistical significance was evaluated with one-way ANOVA and Holm-Sidak post-hoc test, *** p<0.001.

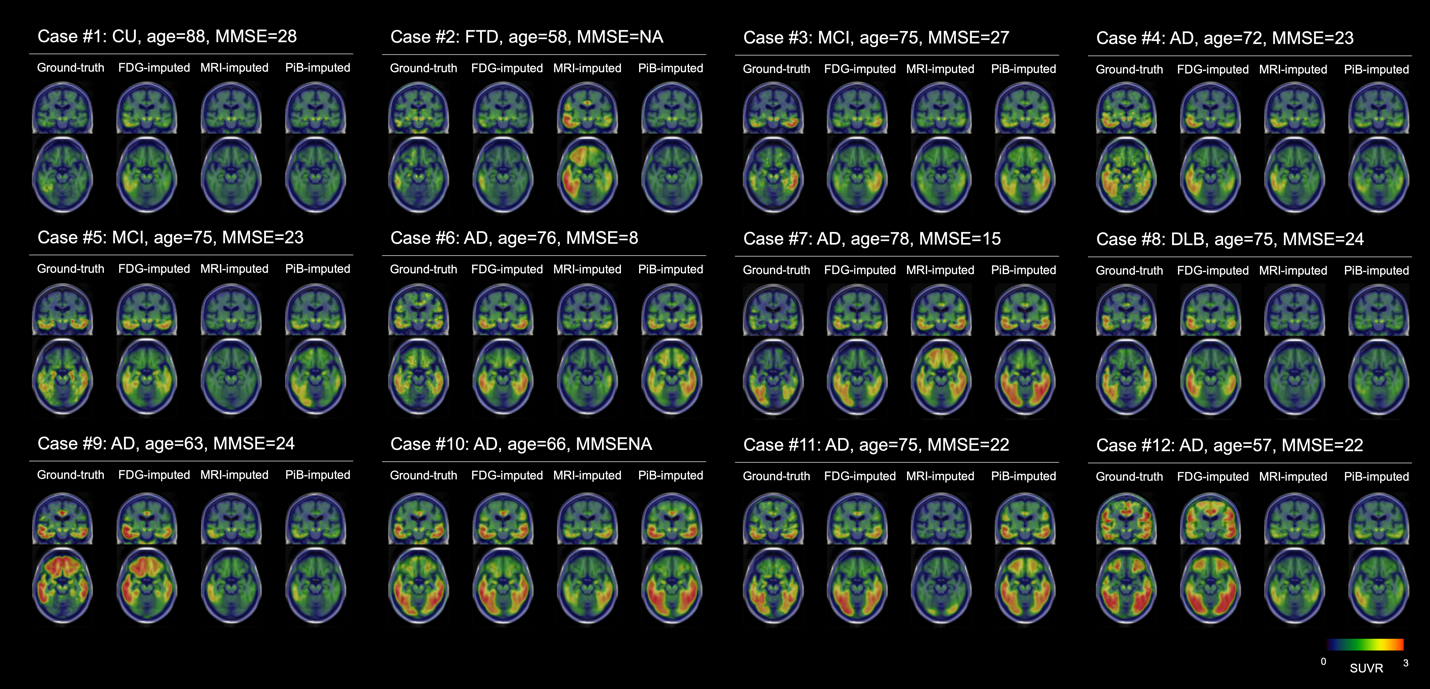

**Supplementary figure 6. Comparisons of performance among the AI-imputed tau PET and ground truth tau PET in SliceView.**

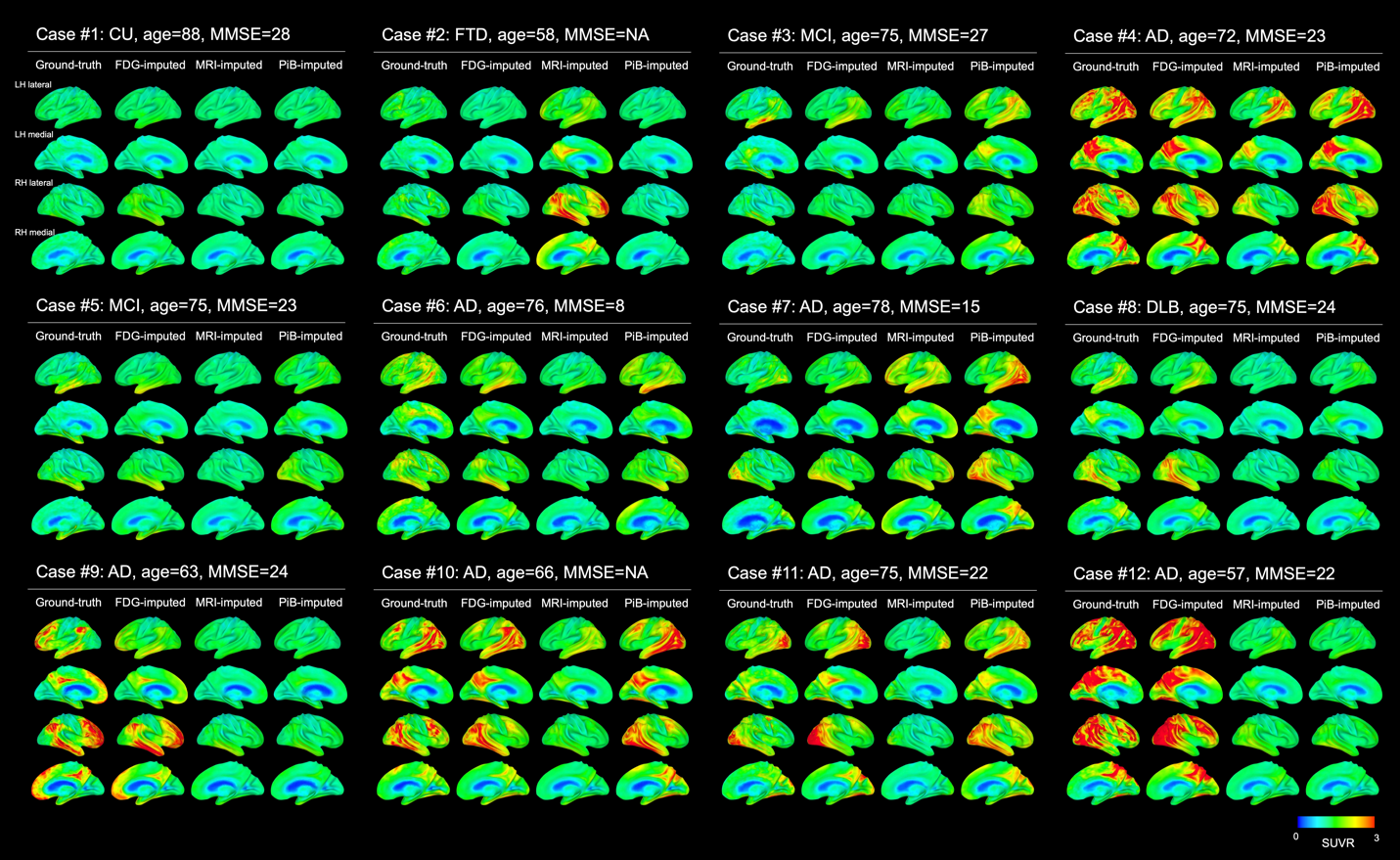

**Supplementary figure 7. 3-dimensional stereotactic surface projection images.**

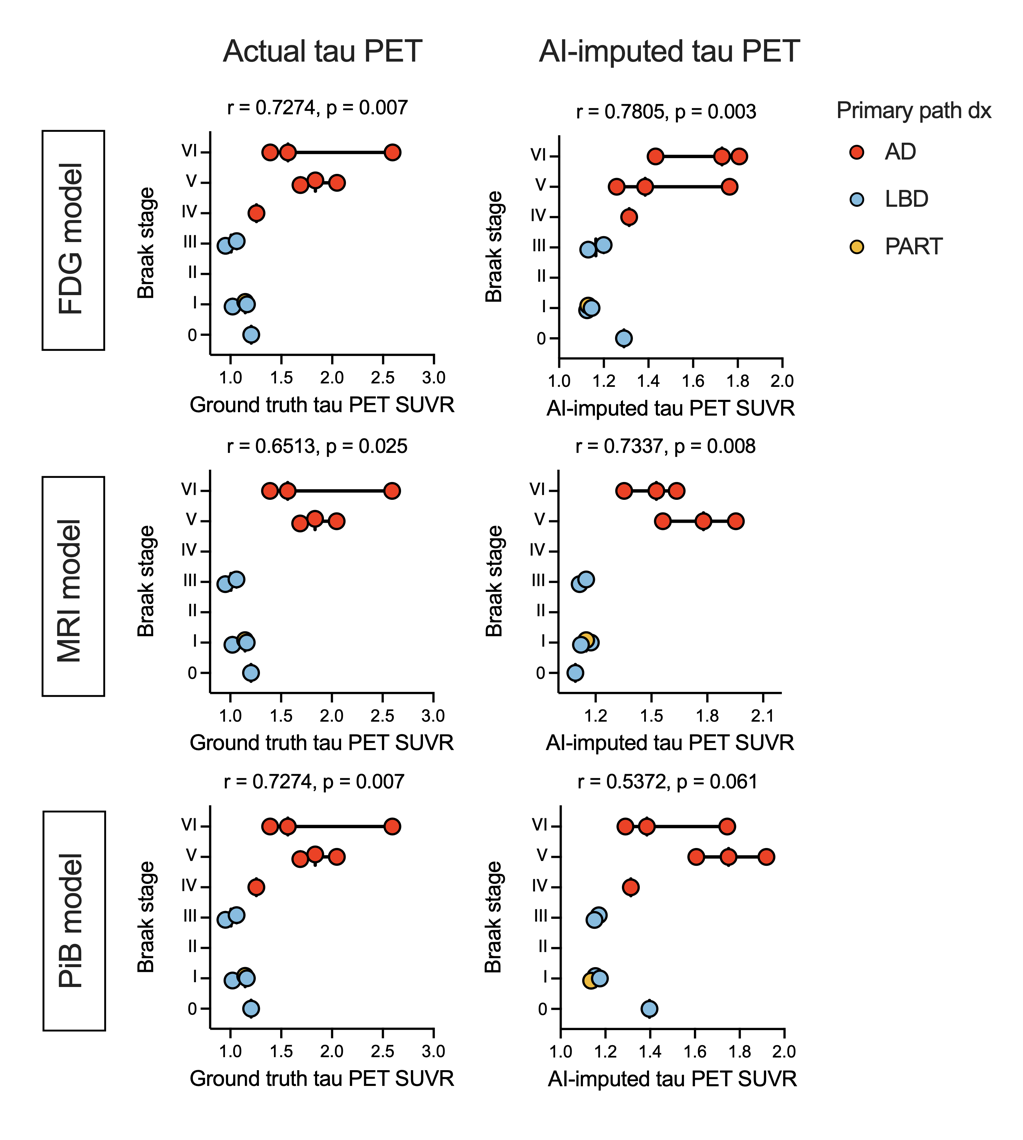

**Supplementary figure 8. AI-imputed tau PET and Braak stage.** Tau PET SUVR using the meta-ROI is shown on the x-axis relative to the Braak stage on the y-axis for the actual tau PET and AI-imputed tau PET. Each participant’s primary pathological diagnosis is visualized as different colors of dots. r, Spearman’s correlation coefficient; p, correlation test p-value. Abbreviations: AD, Alzheimer’s disease; LBD, Lewy body disorders; PART, primary age-related tauopathy.

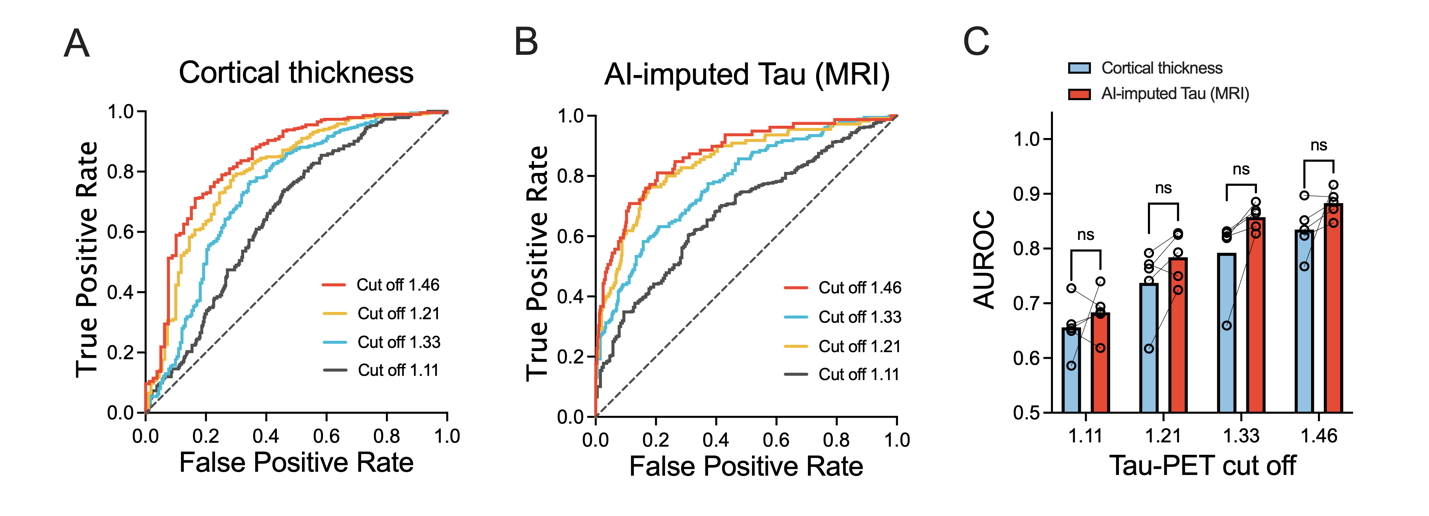

**Supplementary figure 9. ROC analysis for the Siemens cohort.** For cortical thickness, the participants who had Siemens MRI scans were separately analyzed. (A) Cortical thickness (B) MRI-based AI-imputed tau PET (C) AUROC comparison between the cortical thickness and MRI-based AI-imputed tau PET. Statistical significance was assessed with two-way ANOVA and Holm-Sidak post hoc comparison.

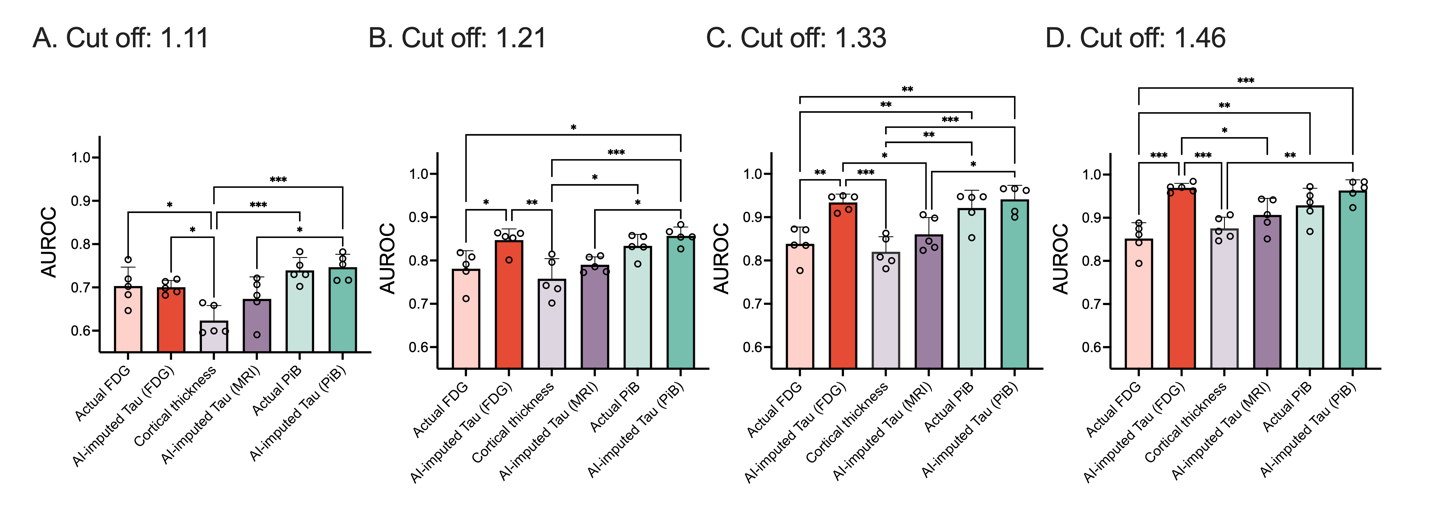

**Supplementary figure 10. AUROC comparisons for tau positivity.** The prediction accuracies (AUROC) for tau positivity were compared among six predictors for (A) 1.11, (B) 1.21, (C) 1.33, and (D) 1.46 cutoff values. Statistical significance was assessed with one-way ANOVA and Holm-Sidak post hoc comparison. Error bars indicate standard deviation. * p<0.05, **p<0.005, *** p<0.001.

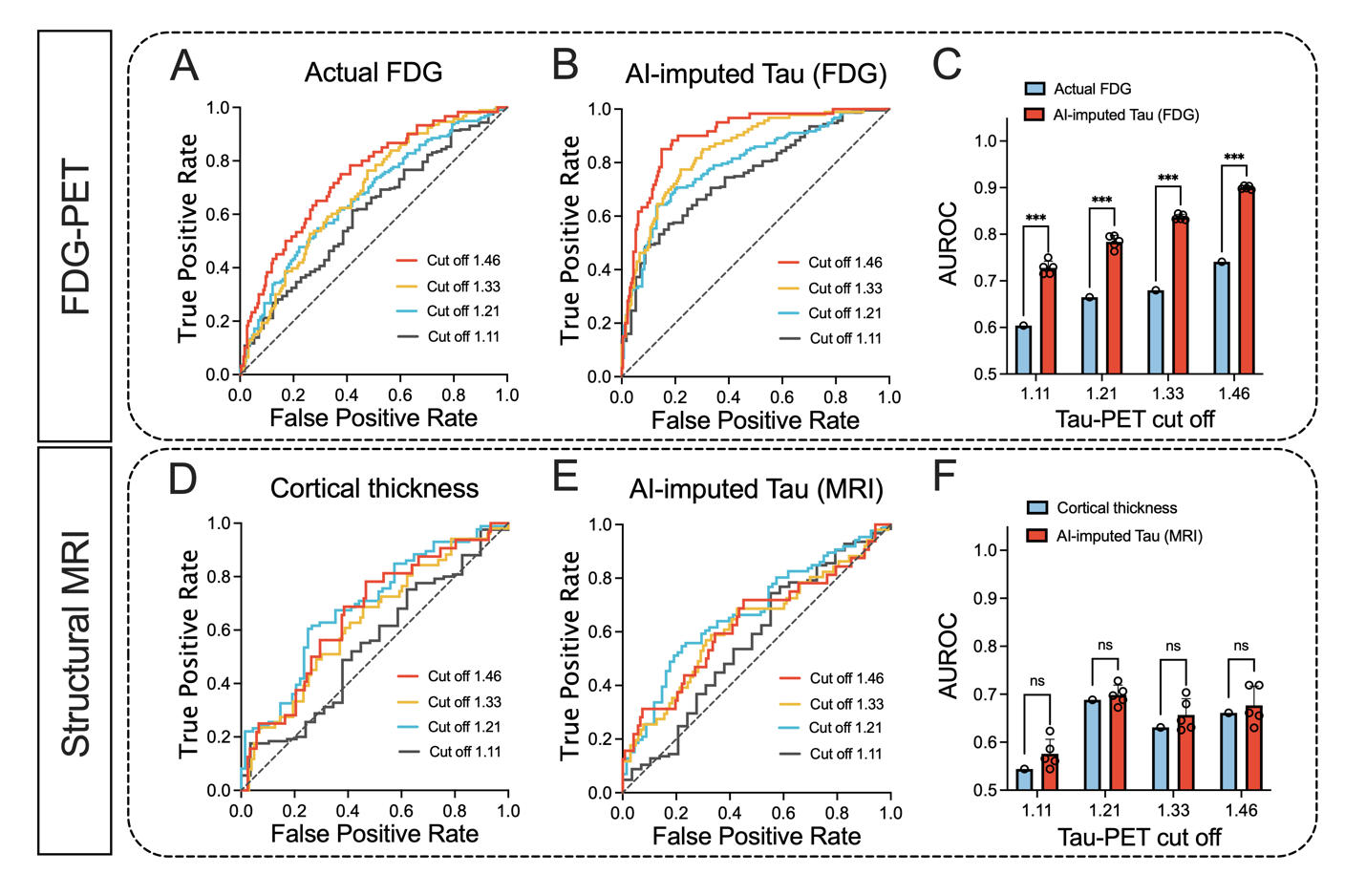

**Supplementary figure 11. ROC curves showing the classification performance on the tau positivity tested on ADNI dataset.** The tau positivity obtained from the ground-truth tau PET using four different meta-ROI cutoff thresholds (1.11, 1.21, 1.33, and 1.46) were predicted. (A) Actual FDG-PET (B) FDG-based AI-imputed tau PET (C) AUROC comparison between the original FDG and FDG-based AI-imputed tau PET. (D) Cortical thickness from the cohort who had Siemens scans (E) MRI-based AI-imputed tau PET from the cohort who had Siemens scans (F) AUROC comparison between the cortical thickness and MRI-based AI-imputed tau PET. Statistical significance was assessed with two-way ANOVA and Holm-Sidak post hoc comparisons. *** p<0.001.

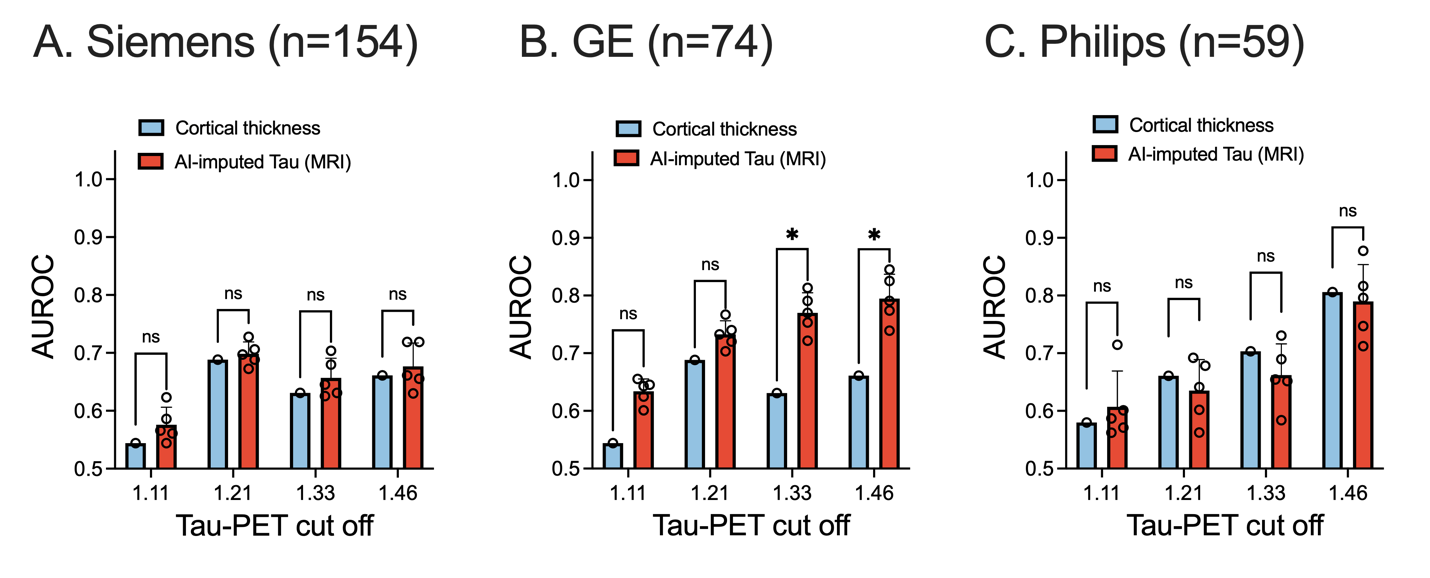

**Supplementary figure 12. AUROC comparisons for different MRI manufacturers tested on ADNI dataset.** The prediction accuracy (AUROC) using four different meta-ROI cutoff thresholds (1.11, 1.21, 1.33, and 1.46) for different MRI manufacturers: (A) The Siemens, (B) GE, and (C) Philips. Statistical significance was assessed with two-way ANOVA and Holm-Sidak post hoc comparisons. * p<0.05.

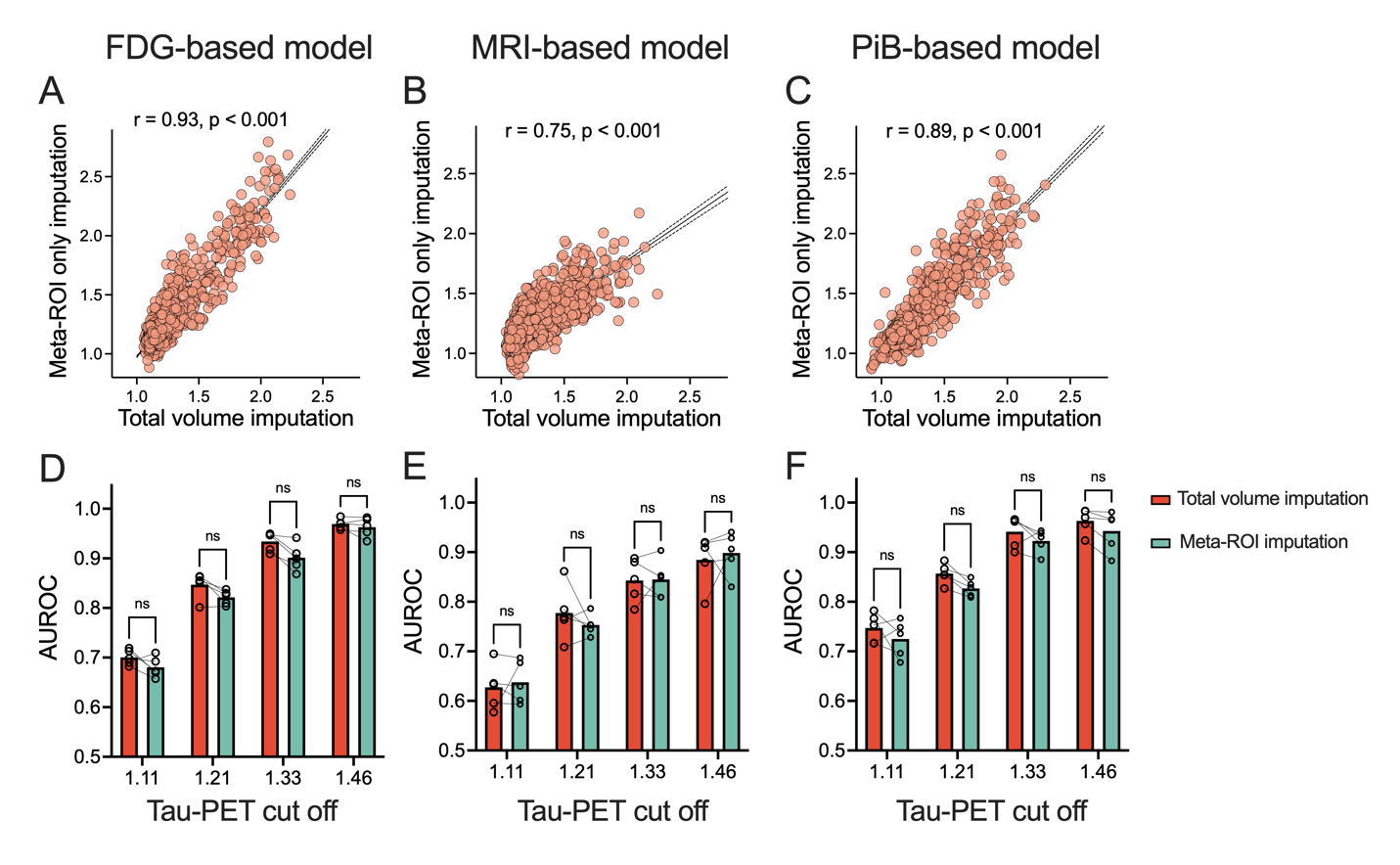

**Supplementary figure 13. Comparison between the total volume imputation vs. meta-ROI only imputation.** (A-C) Scatter plots between the total volume and meta-ROI model. r indicates Pearson’s correlation. (D-F) ROC analysis for tau positivity for FDG-, MRI-, and PiB-based model, respectively. Statistical test was performed using Holm-Sidak test after two-way ANOVA.

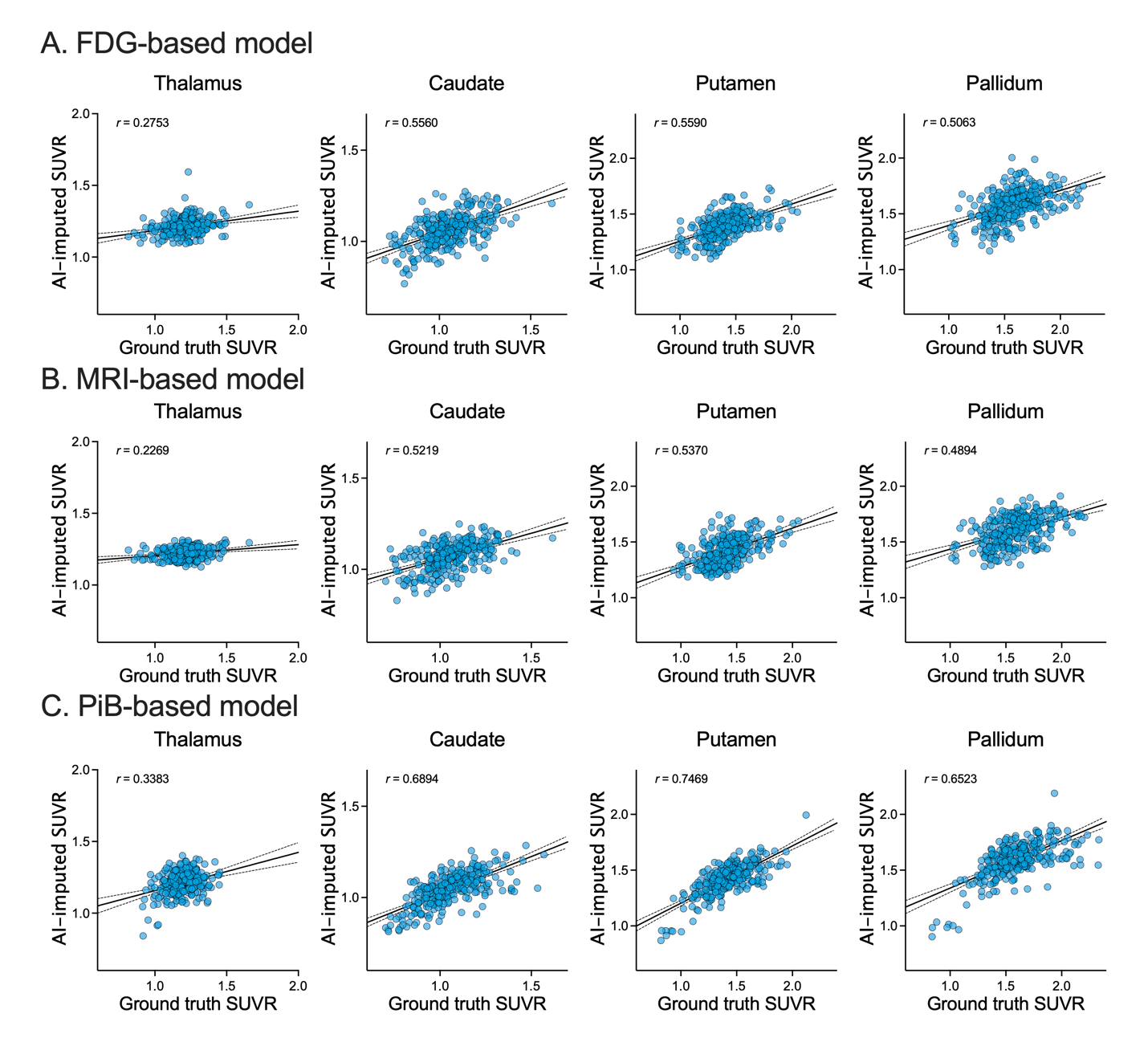

**Supplementary figure 14. AI-imputed SUVR of regions of off-target bindings.** r indicates Pearson’s correlation.

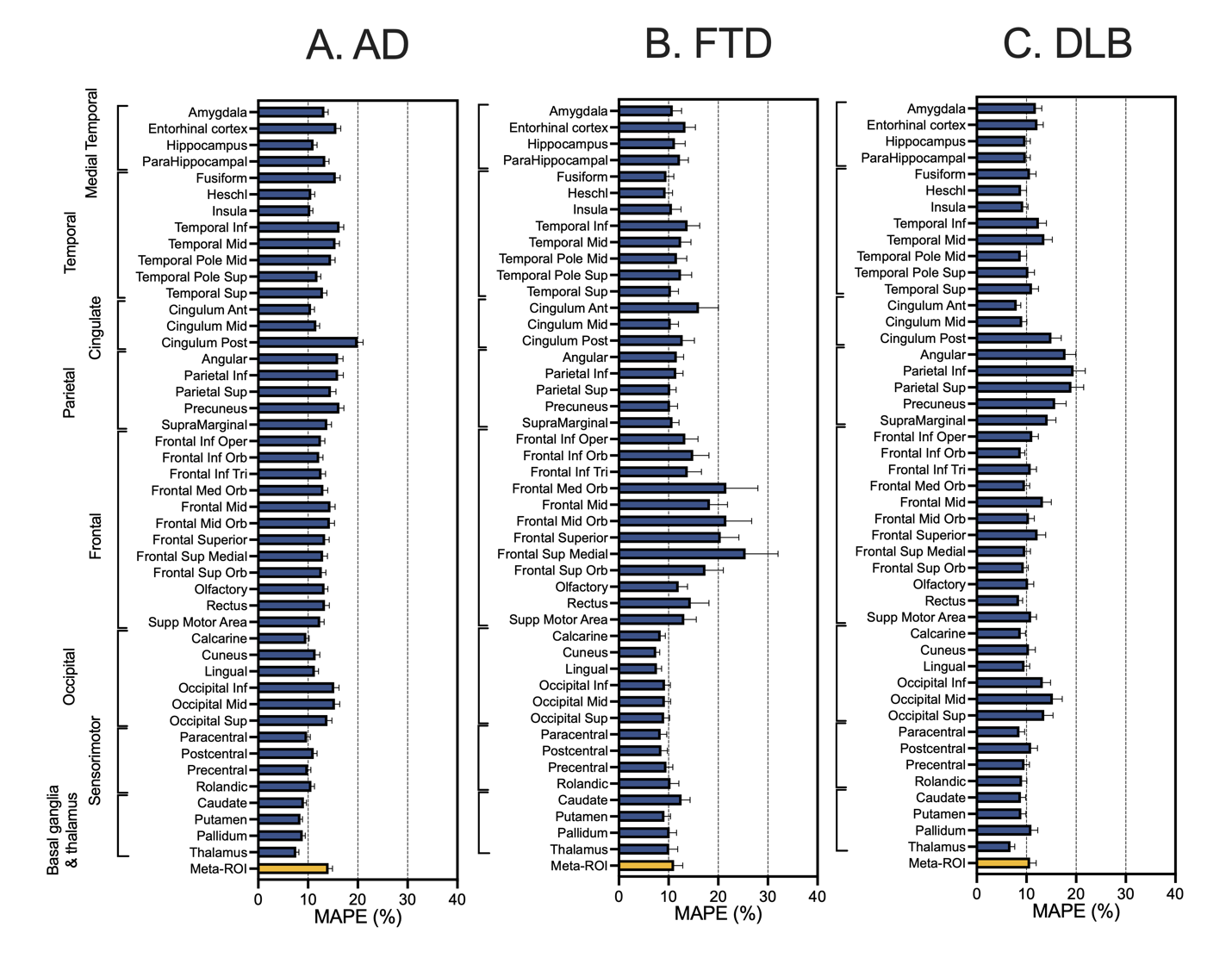

**Supplementary figure 15. Regional MAPE distribution for FDG-based AI-imputed tau PET for AD, FTD, and DLB diagnostic groups.**

**Supplementary Table 1. Demographics for FDG/MRI model.**

Statistical test was performed within each fold. ^1^Linear Model ANOVA, ^2^Pearson’s Chi-squared test, ^3^Median test. Abbreviations: CU: Clinically Unimpaired; MCI: Mild Cognitive Impairment; AD: Alzheimer’s Dementia; DLB: Dementia with Lewy Bodies; FTD: Frontotemporal Dementia; CDR global: Clinical Dementia Rating global; CDR-SB: Clinical Dementia Rating Sum of Boxes; STMS: Short Test of Mental Status; MMSE: Mini-Mental State Examinations.

|  | | **Fold 1** | | | | **Fold 2** | | | | **Fold 3** | | | | **Fold 4** | | | | **Fold 5** | | | |
| --- | --- | --- | --- | --- | --- | --- | --- | --- | --- | --- | --- | --- | --- | --- | --- | --- | --- | --- | --- | --- | --- |
|  | | **Train**  (N=903) | **Validation**  (N=294) | **Test**  (N=308) | **P value** | **Train**  (N=868) | **Validation**  (N=343) | **Test**  (N=294) | **P value** | **Train**  (N=876) | **Validation**  (N=286) | **Test**  (N=343) | **P value** | **Train**  (N=945) | **Validation**  (N=274) | **Test**  (N=286) | **P value** | **Train**  (N=923) | **Validation**  (N=308) | **Test**  (N=274) | **P value** |
| **Age**^1^  Mean (SD) | | 67 (13) | 69 (11) | 67 (12) | 0.387 | 68 (12) | 67 (13) | 69 (11) | 0.116 | 68 (12) | 68 (12) | 67 (13) | 0.163 | 67 (12) | 68 (13) | 68 (12) | 0.537 | 68 (12) | 67 (12) | 68 (13) | 0.558 |
| **Male**^2^  N (%) | | 511 (57%) | 160 (54%) | 183 (59%) | 0.461 | 483 (56%) | 211 (62%) | 160 (54%) | 0.119 | 485 (55%) | 158 (55%) | 211 (62%) | 0.127 | 554 (59%) | 142 (52%) | 158 (55%) | 0.115 | 529 (57%) | 183 (59%) | 142 (52%) | 0.156 |
| **APOE e4 carrier**^2^  N (%)  {N-Miss} | | 303 (35%)  {49} | 110 (39%)  {12} | 97 (32%) {7} | 0.232 | 282 (34%)  {38} | 118 (36%) {18} | 110 (39%) {12} | 0.078 | 297 (35%) {34} | 95 (35%) {16} | 118 (36%) {18} | 0.94 | 325 (36%) {37} | 90 (35%) {15} | 95 (35%) {16} | 0.947 | 323 (37%) {46} | 97 (32%)  {7} | 90 (35%)  {15} | 0.341 |
| **Diagnosis**^2^  N (%) | |  |  |  | **0.007** |  |  |  | **0.004** |  |  |  | 0.209 |  |  |  | 0.207 |  |  |  | 0.441 |
|  | CU | 535 (59%) | 169 (57%) | 186 (60%) |  | 521 (60%) | 200 (58%) | 169 (57%) |  | 513 (59%) | 177 (62%) | 200 (58%) |  | 555 (59%) | 158 (58%) | 177 (62%) |  | 546 (59%) | 186 (60%) | 158 (58%) |  |
|  | MCI | 132 (15%) | 36 (12%) | 40 (13%) |  | 114 (13%) | 58 (17%) | 36 (12%) |  | 113 (13%) | 37 (13%) | 58 (17%) |  | 134 (14%) | 37 (14%) | 37 (13%) |  | 131 (14%) | 40 (13%) | 37 (14%) |  |
|  | AD-spec. | 109 (12%) | 52 (18%) | 33 (11%) |  | 97 (11%) | 45 (13%) | 52 (18%) |  | 122 (14%) | 27 (9%) | 45 (13%) |  | 130 (14%) | 37 (14%) | 27 (9%) |  | 124 (13%) | 33 (11%) | 37 (14%) |  |
|  | DLB-spec. | 52 (6%) | 9 (3%) | 13 (4%) |  | 51 (6%) | 14 (4%) | 9 (3%) |  | 40 (5%) | 20 (7%) | 14 (4%) |  | 36 (4%) | 18 (7%) | 20 (7%) |  | 43 (5%) | 13 (4%) | 18 (7%) |  |
|  | FTD-spec. | 32 (4%) | 3 (1%) | 14 (5%) |  | 34 (4%) | 12 (3%) | 3 (1%) |  | 29 (3%) | 8 (3%) | 12 (3%) |  | 29 (3%) | 12 (4%) | 8 (3%) |  | 23 (2%) | 14 (5%) | 12 (4%) |  |
|  | Other | 43 (5%) | 25 (9%) | 22 (7%) |  | 51 (6%) | 14 (4%) | 25 (9%) |  | 59 (7%) | 17 (6%) | 14 (4%) |  | 61 (6%) | 12 (4%) | 17 (6%) |  | 56 (6%) | 22 (7%) | 12 (4%) |  |
| **CDR global^1^**  Mean (SD)  {N-Miss} | | 0 (0)  {7} | 0 (0)  {5} | 0 (0)  {3} | 0.083 | 0 (0)  {9} | 0 (0)  {1} | 0 (0)  {5} | 0.152 | 0 (0)  {12} | 0 (0)  {2} | 0 (0)  {1} | 0.138 | 0 (0)  {9} | 0 (0)  {4} | 0 (0)  {2} | 0.063 | 0 (0)  {8} | 0 (0)  {3} | 0 (0)  {4} | 0.096 |
| **CDR-SB^1^**  Mean (SD)  {N-Miss} | | 1 (2)  {7} | 2 (3)  {5} | 1 (2)  {3} | 0.055 | 1 (2)  {9} | 1 (2)  {1} | 2 (3)  {5} | 0.123 | 1 (2)  {12} | 1 (2)  {2 } | 1 (2)  {1} | 0.230 | 1 (2)  {9} | 1 (3)  {4} | 1 (2)  {2} | 0.121 | 1 (2)  {8} | 1 (2)  {3} | 1 (3)  {4} | 0.097 |
| **STMS**^1^  Mean (SD)  {N-Miss} | | 33 (6)  {44} | 33 (6)  {20} | 34 (5)  {20} | 0.437 | 33 (6)  {48} | 33 (6)  {16} | 33 (6)  {20} | 0.884 | 33 (6)  {52} | 34 (5)  {16} | 33 (6)  {16} | 0.683 | 33 (6)  {56} | 33 (6)  {12} | 34 (5)  {16} | 0.223 | 33 (6)  {52} | 34 (5)  {20} | 33 (6)  {12} | 0.166 |
| **MMSE**1  Mean (SD)  {N-Miss} | | 27 (4)  {44} | 27 (4)  {20} | 27 (3)  {20} | 0.419 | 27 (4)  {48} | 27 (4)  {16} | 27 (4)  {20} | 0.765 | 27 (4)  {52} | 27 (3)  {16} | 27 (4)  {16} | 0.542 | 27 (4)  {56} | 27 (4)  {12} | 27 (3)  {16} | 0.213 | 27 (4)  {52} | 27 (3)  {20} | 27 (4)  {12} | 0.185 |
| **Tau PET meta-ROI SUVR**^3^  Median  (Q1, Q3) | | 1.21  (1.15, 1.31) | 1.22  (1.15, 1.34) | 1.20  (1.15, 1.29) | 0.065 | 1.21  (1.15, 1.30) | 1.22  (1.14, 1.34) | 1.22  (1.15, 1.34) | 0.259 | 1.21  (1.15, 1.31) | 1.20  (1.13, 1.28) | 1.22  (1.14, 1.34) | 0.303 | 1.21  (1.15, 1.32) | 1.22  (1.17, 1.31) | 1.20  (1.13, 1.28) | 0.124 | 1.21  (1.14, 1.32) | 1.20  (1.15, 1.29) | 1.22  (1.17, 1.31) | **0.034** |

**Supplementary Table 2. Demographics for PiB model**

Statistical test was performed within each fold. ^1^Linear Model ANOVA, ^2^Pearson’s Chi-squared test, ^3^Median test. Abbreviations: CU: Clinically Unimpaired; MCI: Mild Cognitive Impairment; AD: Alzheimer’s Dementia; DLB: Dementia with Lewy Bodies; FTD: Frontotemporal Dementia; CDR global: Clinical Dementia Rating global; CDR-SB: Clinical Dementia Rating Sum of Boxes; STMS: Short Test of Mental Status; MMSE: Mini-Mental State Examinations.

|  | | **Fold 1** | | | | **Fold 2** | | | | **Fold 3** | | | | **Fold 4** | | | | **Fold 5** | | | |
| --- | --- | --- | --- | --- | --- | --- | --- | --- | --- | --- | --- | --- | --- | --- | --- | --- | --- | --- | --- | --- | --- |
|  | | **Train**  (N=885) | **Validation**  (N=301) | **Test**  (N=293) | **P value** | **Train**  (N=859) | **Validation**  (N=319) | **Test**  (N=301) | **P value** | **Train**  (N=876) | **Validation**  (N=284) | **Test**  (N=319) | **P value** | **Train**  (N=913) | **Validation**  (N=282) | **Test**  (N=284) | **P value** | **Train**  (N=904) | **Validation**  (N=293) | **Test**  (N=282) | **P value** |
| **Age**^1^  Mean (SD) | | 67 (12) | 68 (12) | 69 (12) | 0.323 | 67 (12) | 69 (12) | 68 (12) | 0.210 | 68 (12) | 66 (12) | 69 (12) | 0.02 | 68 (12) | 67 (13) | 66 (12) | 0.012 | 68 (12) | 69 (12) | 67 (13) | 0.378 |
| **Male**^2^  N (%) | | 496 (56%) | 171 (57%) | 173 (59%) | 0.668 | 486 (57%) | 183 (57%) | 171 (57%) | 0.971 | 493 (56%) | 164 (58%) | 183 (57%) | 0.886 | 527 (58%) | 149 (53%) | 164 (58%) | 0.329 | 518 (57%) | 173 (59%) | 149 (53%) | 0.287 |
| **APOE e4 carrier**^2^  N (%)  {N-Miss} | | 303 (36%)  {46} | 105 (36%)  {9} | 96 (34%)  {13} | 0.854 | 288 (35%)  {44} | 111 (37%)  {15} | 105 (36%)  {9} | 0.931 | 293 (35%)  {35} | 100 (38%)  {18} | 111 (37%)  {15} | 0.679 | 312 (36%)  {37} | 92 (34%)  {13} | 100 (38%)  {18} | 0.711 | 316 (37%)  {42} | 96 (34%)  {13} | 92 (34%)  {13} | 0.653 |
| **Diagnosis**^2^  N (%) | |  |  |  | **0.008** |  |  |  | **0.018** |  |  |  | 0.333 |  |  |  | **0.026** |  |  |  | **0.017** |
|  | CU | 538 (61%) | 166 (55%) | 183 (62%) |  | 544 (63%) | 177 (55%) | 166 (55%) |  | 531 (61%) | 179 (63%) | 177 (55%) |  | 526 (58%) | 182 (65%) | 179 (63%) |  | 522 (58%) | 183 (62%) | 182 (65%) |  |
|  | MCI | 112 (13%) | 42 (14%) | 50 (17%) |  | 112 (13%) | 50 (16%) | 42 (14%) |  | 124 (14%) | 30 (11%) | 50 (16%) |  | 142 (16%) | 32 (11%) | 30 (11%) |  | 122 (13%) | 50 (17%) | 32 (11%) |  |
|  | AD-spec. | 125 (14%) | 38 (13%) | 27 (9%) |  | 111 (13%) | 41 (13%) | 38 (13%) |  | 107 (12%) | 42 (15%) | 41 (13%) |  | 106 (12%) | 42 (15%) | 42 (15%) |  | 121 (13%) | 27 (9%) | 42 (15%) |  |
|  | DLB-spec. | 40 (5%) | 25 (8%) | 7 (2%) |  | 29 (3%) | 18 (6%) | 25 (8%) |  | 41 (5%) | 13 (5%) | 18 (6%) |  | 50 (5%) | 9 (3%) | 13 (5%) |  | 56 (6%) | 7 (2%) | 9 (3%) |  |
|  | FTD-spec. | 18 (2%) | 12 (4%) | 10 (3%) |  | 18 (2%) | 10 (3%) | 12 (4%) |  | 27 (3%) | 3 (1%) | 10 (3%) |  | 32 (4%) | 5 (2%) | 3 (1%) |  | 25 (3%) | 10 (3%) | 5 (2%) |  |
|  | Other | 52 (6%) | 18 (6%) | 16 (5%) |  | 45 (5%) | 23 (7%) | 18 (6%) |  | 46 (5%) | 17 (6%) | 23 (7%) |  | 57 (6%) | 12 (4%) | 17 (6%) |  | 58 (6%) | 16 (5%) | 12 (4%) |  |
| **CDR global^1^**  Mean (SD)  {N-Miss} | | 0 (0)  {4} | 0 (0)  {3} | 0 (0)  {8} | **0.017** | 0 (0)  {12} | 0 (0)  {0} | 0 (0)  {3} | **0.009** | 0 (0)  {11} | 0 (0)  {4} | 0 (0)  {0} | 0.610 | 0 (0)  {11} | 0 (0)  {0} | 0 (0)  {4} | 0.337 | 0 (0)  {7} | 0 (0)  {8} | 0 (0)  {0} | 0.087 |
| **CDR-SB^1^**  Mean (SD)  {N-Miss} | | 1 (2)  {4} | 2 (3)  {3} | 1 (2)  {8} | **0.021** | 1 (2)  {12} | 1 (2)  {0} | 2 (3)  {3} | **0.007** | 1 (2)  {11} | 1 (2)  {4} | 1 (2)  {0} | 0.351 | 1 (2)  {11} | 1 (2)  {0} | 1 (2)  {4} | 0.320 | 1 (2)  {7} | 1 (2)  {8} | 1 (2)  {0} | 0.053 |
| **STMS**^1^  Mean (SD)  {N-Miss} | | 33 (6)  {37} | 33 (7)  {23} | 34 (5)  {15} | 0.054 | 34 (5)  {41} | 33 (6)  {11} | 33 (7)  {23} | **0.023** | 33 (6)  {44} | 34 (5)  {20} | 33 (6)  {11} | 0.160 | 33 (6)  {49} | 33 (6)  {6} | 34 (5)  {20} | 0.167 | 33 (6)  {54} | 34 (5)  {15} | 33 (6)  {6} | 0.329 |
| **MMSE**1  Mean (SD)  {N-Miss} | | 27 (4)  {37} | 27 (4)  {23} | 27 (3)  {15} | 0.058 | 27 (3)  {41} | 27 (4)  {11} | 27 (4)  {23} | **0.024** | 27 (4)  {44} | 27 (3)  {20} | 27 (4)  {11} | 0.152 | 27 (4)  {49} | 27 (4)  {6} | 27 (3)  {20} | 0.170 | 27 (4)  {54} | 27 (3)  {15} | 27 (4)  {6} | 0.300 |
| **Tau PET meta-ROI SUVR**^3^  Median  (Q1, Q3) | | 1.21  (1.15, 1.31) | 1.21  (1.15, 1.33) | 1.21  (1.15, 1.30) | 0.770 | 1.21  (1.14, 1.30) | 1.22  (1.15, 1.34) | 1.21  (1.15, 1.33) | 0.806 | 1.21  (1.15, 1.31) | 1.21  (1.14, 1.30) | 1.22  (1.15, 1.34) | 0.830 | 1.21  (1.15, 1.32) | 1.21  (1.14, 1.30) | 1.21  (1.14, 1.30) | 0.912 | 1.21  (1.15, 1.32) | 1.21  (1.15, 1.30) | 1.21  (1.14, 1.30) | 0.663 |

**Supplementary Table 3. ADNI Cohort Demographics**

| Characteristic | Clinical Diagnosis | | |
| --- | --- | --- | --- |
|  | Normal | MCI | Dementia |
| N (%) | 15 (5.21) | 205 (71.18) | 68 (23.61) |
| Age, median (min max), years | 68 (56 89) | 75 (56 92) | 77.5 (56 92) |
| Male sex, n (%) | 7 (46.67) | 117 (57.07) | 39 (57.35) |
| Education, median (IQR), years | 16 (14 18) | 16 (14 18) | 16 (13.5 18) |
| Clinical Dementia Rating Scale-Sum of Boxes, median (IQR) | 0 (0 0) | 1 (0.5 2) | 4.5 (3.5 5.5) |
| Meta-ROI FDG PET SUVR, median (IQR) | 1.27 (1.24 1.29) | 1.20 (1.13 1.27) | 1.10 (1.03 1.17) |
| Meta-ROI Tau PET SUVR,  median (IQR) | 1.12 (1.06 1.15) | 1.20 (1.12 1.33) | 1.45 (1.26 1.67) |
